# *AlleleShift*: An R package to predict and visualize population-level changes in allele frequencies in response to climate change

**DOI:** 10.1101/2021.01.15.426775

**Authors:** Roeland Kindt

## Abstract

**Background:** At any particular location, frequencies of alleles in organisms that are associated with adaptive traits are expected to change in future climates through local adaption and migration, including assisted migration (human-implemented when climate change is more rapid than natural migration rates). Making the assumption that the baseline frequencies of alleles across environmental gradients can act as a predictor of patterns in changed climates (typically future but possibly paleo-climates), a methodology is provided by *AlleleShift* of predicting changes in allele frequencies at populations’ locations.

**Methods:** The prediction procedure involves a first calibration and prediction step through redundancy analysis (RDA), and a second calibration and prediction step through a generalized additive model (GAM) with a binomial family. As such, the procedure is fundamentally different to an alternative approach recently proposed to predict changes in allele frequencies from canonical correspondence analysis (CCA). My methodology of *AlleleShift* is also different in modelling and predicting allele counts through constrained ordination (not frequencies as in the CCA approach) and modelling both alleles for a locus (not solely the minor allele as in the CCA method; both methods were developed for diploid organisms where individuals are homozygous (AA or BB) or heterozygous (AB)). Whereas the GAM step ensures that allele frequencies are in the range of 0 to 1 (negative values are sometimes predicted by the RDA and CCA approaches), the RDA step is based on the Euclidean distance that is also the typical distance used in Analysis of Molecular Variance (AMOVA). The *AlleleShift::amova.rda* enables users to verify that the same ‘mean-square’ values are calculated by AMOVA and RDA, and gives the same final statistics with balanced data.

**Results:** Besides data sets with predicted frequencies, *AlleleShift* provides several visualization methods to depict the predicted shifts in allele frequencies from baseline to changed climates. These include ‘dot plot’ graphics (function *shift.dot.ggplot*), pie diagrams (*shift.pie.ggplot*), moon diagrams (*shift.moon.ggplot*), ‘waffle’ diagrams (*shift.waffle.ggplot*) and smoothed surface diagrams of allele frequencies of baseline or future patterns in geographical space (*shift.surf.ggplot*). As these were generated through the *ggplot2* package, methods of generating animations for a climate change time series are straightforward, as shown in the documentation of *AlleleShift* and in the supplementary materials. In addition, graphical methods are provided of showing shifts of populations in environmental space (*population.shift*) and to assess how well the predicted frequencies reflect the original frequencies for the baseline climate (*freq.ggplot*).

**Availability:** *AlleleShift* is available as an open-source R package from https://github.com/RoelandKindt/AlleleShift. Genetic input data is expected to be in the *adegenet::genpop* format, which can be generated from the *adegenet::genind* format. Climate data is available from various resources such as *WorldClim* and *Envirem*.

## Introduction

There is clear evidence of climate change, with our planet facing a climate emergency (Ripple *et al.*, 2020). Anticipating climate change, many countries are developing National Adaptation Plans (NAPs). Specifically for forests and trees, technical guidelines have recently been prepared on the integration and forests, agroforestry and trees in the formulation and implementations of NAPs (Meybeck et al., 2020). Stanturf et al. (2015) provide a practical framework and a stoplight tool to plan for climate change mitigation and adaptation in forest and landscape restoration initiatives.

For loci involved in adaptation, shifts in allele frequencies (and changes in phenotypes as a result) can be anticipated (Günther & Coop, 2013; Stange, Barrett & Hendry, 2020). Although the methods involved are far from straightforward, statistical approaches such as Genome Wide Association Studies (GWAS) and Environmental Association Analyses (EAA) can be applied to genomic data to postulate genes and specific alleles involved in climate change responses (Luikart et al., 2018; Anderson & Song, 2020; Waldvogel et al. 2020).

The methodology used in *AlleleShift* to predict changes in allele frequencies due to climate change was inspired by a protocol developed by Blumstein et al. (2020), but is also fundamentally different, as documented in Table 1. Both methods exploit the analogy between analysing a community matrix (consisting of sites as rows, species as columns and abundances as cell values) and a genetic matrix (consisting of populations or individuals as rows, alleles as columns and allele counts as cell values). As a consequence, both rely on common constrained ordination methods in the field of community ecology: redundancy analysis (RDA) or canonical correspondence analysis (CCA). A thorough discussion of both methods is available in Legendre & Legendre (2012, pp. 629 - 673), a key reference work in the field.

**Table 1.** Fundamental differences between the CCA approach proposed by Blumstein et al. (2020) and methods available via AlleleShift

| **Blumstein et al. (2020)** | **AlleleShift** |
| --- | --- |
| Calculate allele frequencies with a custom function ‘freq’ | • Import data in a `adegenet::genind` object<br>• Convert data to `adegenet::genpop` object<br>• Calculate allele frequencies via `adegenet::makefreq`<br>• Calculate population level statistics in `poppr::poppr`<br>• Check correlations among alleles |
| Principal Components Analysis to reduce number of explanatory variables for a constrained ordination analysis | • Variance Inflation Factor analysis to reduce the explanatory variables to a subset with VIF below the threshold of 0.95<br>• Options of pre-selecting (keeping) certain explanatory variables (for example, these could be the three bioclimatic variables selected by Booth (2016) for estimating future ranges of tree species) |
| Combine community and environmental data sets to one data set, but split again during analysis | Keep community and environmental data set separate throughout the analysis, being cautious in not reordering any of the data sets. |
| (not implemented) | • Connect baseline and future positions of populations in ordination diagrams<br>• Fit superellipses and conduct RDA to check for niche differentiation between baseline and future positions of climates |
| Fit constrained ordination model of canonical correspondence analysis (CCA) via `vegan::cca` | Fit constrained ordination model of redundancy analysis (RDA) via `vegan::rda`<br>**See more details in the main text about the justification of using RDA instead of CCA** |
| Fit CCA with the allele frequencies in the community data set | Fit RDA with allele counts in the community data set |
| Fit CCA only with the minor alleles (A) | Fit RDA with all alleles (A and B) |
| Reduce the number of explanatory variables | Possible, but not recommended as this reduces the explanatory power of the model to predict allele frequencies |
| (not implemented) | • Calibrate a Generalized Additive Model (GAM) that ensures predicted allele frequencies are within the range of 0 to 1<br>• Estimate standard errors for predictions |

| Blumstein et al. (2020) | AlleleShift |
| --- | --- |
| (not implemented) | • Show the population identities in graphs of actual allele frequencies vs. predicted frequencies<br>• Show the allele identities in graphs of actual allele frequencies vs. predicted allele frequencies<br>• Split the graph between populations with high $R^2$ and low $R^2$ values |
| (not implemented) | Various visualization methods of shifts in allele frequencies, including animation methods |

One of the fundamental differences I implement compared to Blumstein et al. (2020) is to use RDA instead of CCA.

This is for several reasons:

- RDA is based on Euclidean distances and without explanatory variables is equivalent to principal components analysis (PCA). Euclidean distances are also used in Analysis of Molecular Variance (AMOVA; Excoffier, Smouse & Quattro, 1992; Meirmans & Liu, 2018; Michalakis & Excoffier, 1996) and it can be demonstrated (see examples for the `AlleleShift::amova.rda` function and possibly also compare with AMOVA analysis in GenAlEx; Peakal & Smouse, 2012) that RDA provides the same information on squared Euclidean distances and mean squares as an AMOVA analysis.
- In supplementary materials A, I demonstrate how Euclidean distances between `adegenet::genpop` objects are linearly related to the Euclidean distances between the centroids obtained from a PCA analysis of `adegenet::genind` objects. As a corollary, shifts of populations can be understood as the average shift of individuals in ordination space.
- Recently, I also showed (Kindt 2020a) how RDA can be directly interpreted in terms of Sums-of-Squares of AMOVA by analysing distances from individuals to centroids and among centroids.
- Various recent studies of adaptative genetic variation have also used the RDA methodology (e.g., Razgour et al.; 2019; Capblancq et al., 2020; Nelson, Motamayor & Cornejo, 2020; Temunović et al., 2020).
- Theoretically, RDA is better for analysis of linear patterns in species abundances, whereas CCA is more appropriate for analyzing unimodal patterns For the particular question of whether allele frequencies increase or decrease (i.e., show a linear trend) in future climates, I consider a method that assumes linear patterns is more appropriate. Smoothed regression surfaces of allele frequencies in baseline climates (see Visualizations in the results section) showed that major linear trends in allele frequencies were indeed linear rather than unimodal.
- The interpretation of species scores (here: allele scores) in ordination diagrams generated by RDA is straightforward as showing the general direction of increasing abundances (here: increasing allele counts). The interpretation of species scores in ordination diagrams generated by CCA is more complex in showing the peak of its unimodal distribution against a vector of an explanatory variable (see Fig. 3 in Ter Braak 1987; Fig. 11.9 in Legendre & Legendre, 2012; or Figure 10.13 in Kindt & Coe, 2002).

## Materials & Methods

### Data import

Genetic response data (including a matrix with populations as rows and allele counts as columns) for the calibration of the `AlleleShift::count.model` and prediction via `AlleleShift::count.pred` is required to be in the `adegenet::genpop` format. Individual-based data that are in the `adegenet::genind` format can be converted into the genpop format via the `adegenet::genind2genpop` function. The *adegenet* and *poppr* packages provide various methods of importing data from other software application formats into the *genind* format, such as `adegenet::import2genind` and `poppr::read.genalex`. Environmental data of populations, used as explanatory variables in redundancy analysis (RDA), is expected to be provided as a data.frame with the same sequence of populations as the genetic response data (this is a general requirement for community ecology methods in the *vegan* and *BiodiversityR* packages; a check is available via `BiodiversityR::check.datasets`). Whereas environmental data typically is baseline and changed (bio)climatic data such as is available from WorldClim (Fick & Hijmans, 2017), ENVIREM (Title & Bemmels, 2018) or PaleoClim (Brown *et al.*, 2018), it is also possible to expand input to other data available for species distribution modelling (a recent overview of available data sets is provided by Booth, 2018).

### Data analysis

Prior to calibrating the models that predict allele frequencies from bioclimatic explanatory data, it is recommended to reduce the explanatory variables to a subset where the Variance Inflation Factor (VIF) is below a predefined threshold for each variable. Such methods are also recommended for regression analysis (Fox & Monette, 1992) and species distribution modelling (Kindt, 2018). With this approach, it is easy to select the same variables from future data sets and comparison with other studies may also become easier. VIF analysis, and an additional feature of forcing the algorithm to keep preselected variables within the final subset, is available via `AlleleShift::VIF.subset` (Table 2). This is a function that uses `BiodiversityR::ensemble.VIF.dataframe` internally after a first step of removing all explanatory variables that have correlations larger than the VIF threshold with the preselected variables. There is an option to generate a correlation matrix for the final subset of variables (Figure 1). I also recommend conducting the VIF analysis for the genetic response data (see discussion and Figure 1). I further advise to remove any individuals with partially missing genetic or (bio)climatic data prior to the analysis.

**Table 2.** Functions found in AlleleShift and their short descriptions

| Function | Description |
| --- | --- |
| **Preparation** |  |
| VIF.subset | Reduce the number of explanatory variables through Variance Inflation Factor analysis, with an option to plot a correlation matrix (GGally::ggcorr). Internally, the function calls the BiodiversityR::ensemble.VIF.dataframe function. |
| **Analysis** |  |
| count.model | Calibration of RDA model (vegan::rda) with baseline allele counts as response and baseline bioclimatic variables as explanatory variables |
| count.pred | Prediction of allele counts (vegan::predict.rda) from explanatory variables. Explanatory variables correspond to the baseline climate to check the calibration |
| freq.model | Calibration of GAM model (mgcv::gam) with baseline allele frequencies as response and predicted baseline counts from the RDA model as explanatory variables |
| freq.pred | Prediction of allele frequencies (mgcv::predict.gam) from the predicted alleles counts of count.pred |
| amova.rda | Perform AMOVA with the outputs from RDA. The function returns an output that is similar to the output of poppr::poppr.amova so that results can be readily compared |
| **Visualization** |  |
| population.shift | Shifts of populations in environmental space, with superellipses (ggforce::geom_mark_ellipse) and arrows between baseline and changed positions to show climatic shifts. Internally, the function calls vegan::ordiplot and various helper functions from BiodiversityR (Kindt 2020a) are used |
| freq.ggplot | Plots of baseline allele frequencies against predicted allele frequencies. Data points can be coloured differently by population or by allele |
| shift.dot.ggplot | Shifts of Allele Frequencies as Response to Climate Change |
| shift.pie.ggplot | Shifts of Allele Frequencies as Response to Climate Change via ggforce::geom_arc_bar |
| pie.baker | Helper function to prepare data for shift.pie.ggplot from the output of freq.pred. |
| shift.moon.ggplot | Shifts of Allele Frequencies as Response to Climate Change via ggibbous::geom_moon |
| moon.waxer | Helper function to prepare data for shift.moon.ggplot from the output of freq.pred |
| shift.waffle.ggplot | Shifts of Allele Frequencies as Response to Climate Change. Graphics are similar to waffle::waffle, but the graph is made *de novo* in AlleleShift |
| `waffle.baker` | Helper function to prepare data for `shift.waffle.ggplot` from the output of `freq.pred` |
| `shift.surf.ggplot` | Shifts of Allele Frequencies as Response to Climate Change, plotted in geographical space through smoothed regression surfaces (`vegan::ordisurf`) |

**Figure 1.**
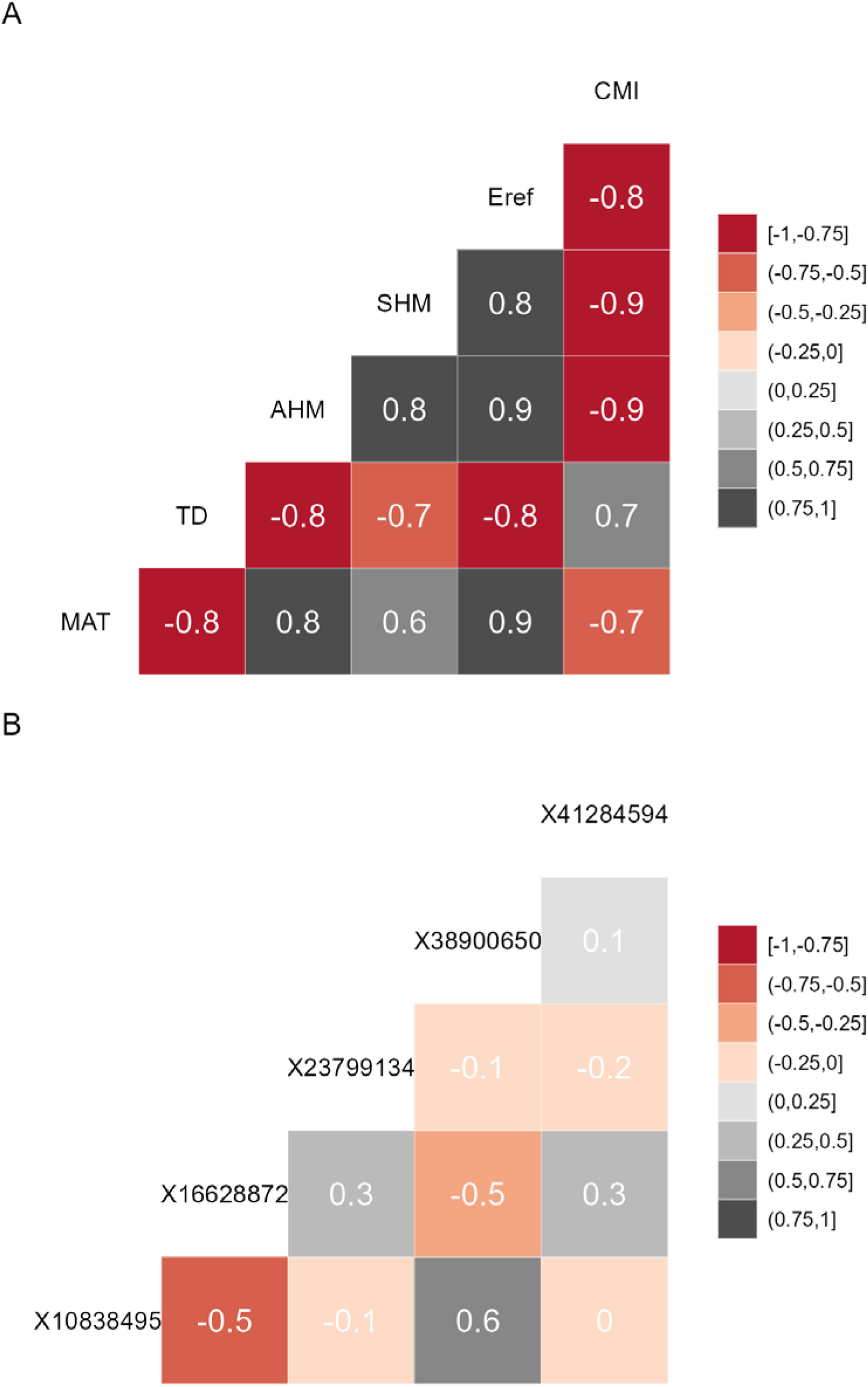
(A) Correlation matrix plot for the final subset of bioclimatic variables selected by the `VIF.subset` function. (B) Correlation matrix plot for the minor alleles.

Prior to model calibration, I suggest checking (according to criteria described below) for the shifts of populations in environmental space between the baseline and changed climates. This can be achieved via function `AlleleShift::population.shift`, which draws arrows between each population in the baseline and changed climate. There are alternatives of using principal components analysis (PCA) or redundancy analysis (RDA) to generate the ordination diagrams (Figure 2). What is important to check in the ordination graphs is whether populations shift in a similar fashion, as that will facilitate the interpretation of predicted shifts in allele frequencies.

**Figure 2.**
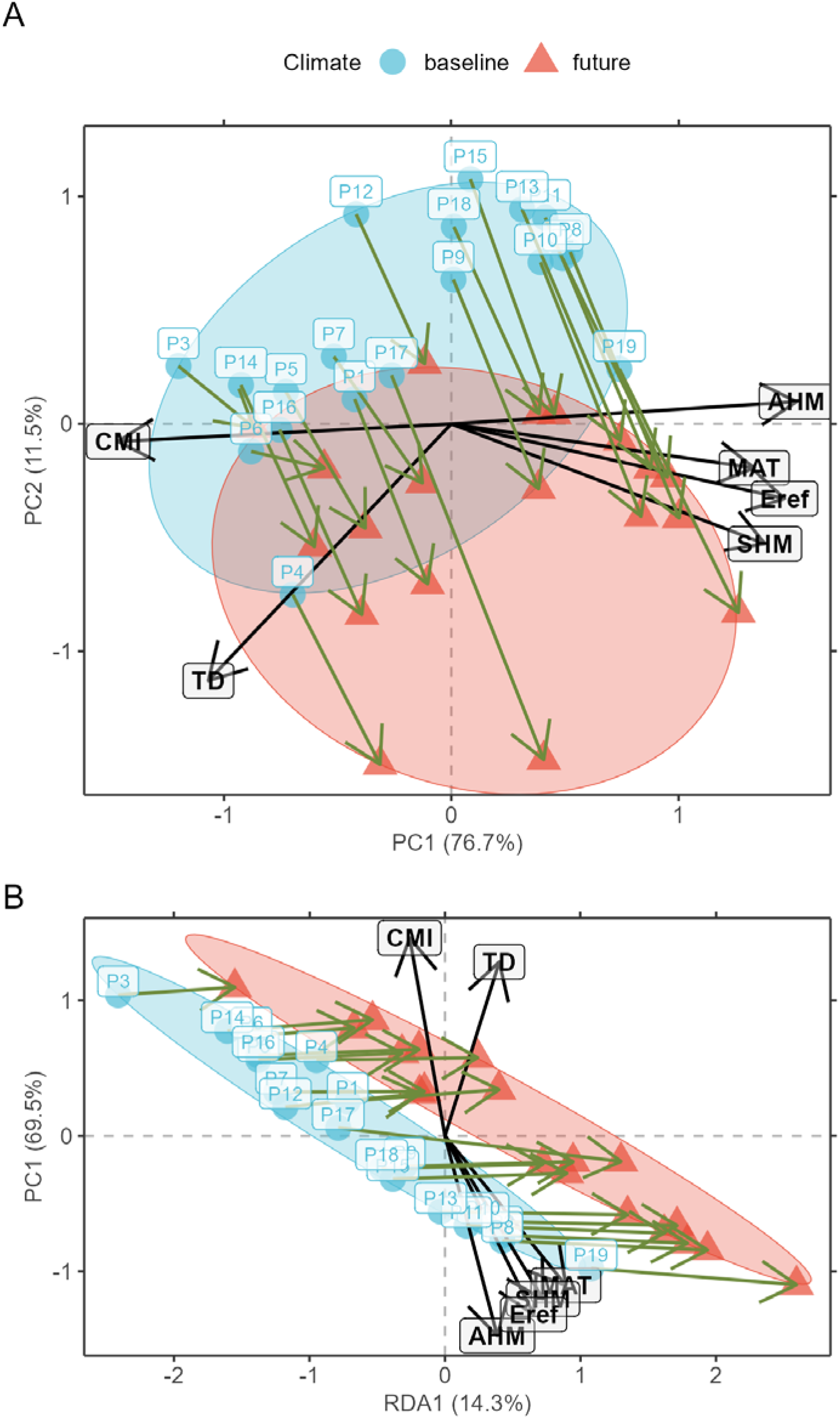
(A) Principal Component Analysis (PCA) ordination graph for shifts in populations in environmental space. (B) Redundancy Analysis (RDA) ordination graph for shifts in populations with climate (a categorical variable with ‘baseline’ and ‘future’ as levels) as explanatory variable.

With the selected genetic and explanatory data, model calibration can be done. In a first step, a RDA model is fitted (`AlleleShift::count.model`) that can predict counts of alleles in baseline or changed climates (`AlleleShift::count.pred`). The user has the option also to obtain results for the canonical correspondence analysis procedure of Blumstein et al. (2020) with the count model via argument `cca.model=TRUE`. In the second step, the predicted allele counts for the baseline climate serve as explanatory variables for the calibration of a generalized linear model (GAM via `mgcv::gam`; Wood, 2004) with the baseline allele frequencies as response and a binomial family function (`AlleleShift::freq.model`). This procedure ensures that predictions are within the realistic interval for frequencies between 0 to 1. Function `AlleleShift::freq.pred` allows the prediction of allele frequencies for baseline and changed climates.

The output of the two steps (RDA followed by GAM) is a data.frame as shown in Table 3 (to fit printing space available here, the data is shown in a transposed format where rows and columns were swapped). All figures shown in this manuscript were obtained with the example data sets of `AlleleShift::Poptri.genind` (individually-based allele counts), `AlleleShift::Poptri.baseline.env` (climatic descriptors of the populations in the baseline climate), `AlleleShift::Poptri.future.env` (climatic descriptors of the populations in the future climate) and `AlleleShift::Poptri.loc` (geographical coordinates of the populations). These data sets were converted from the example data sets provided by Blumstein et al. (2020) for *Populus trichocarpa*. It can be seen for population Nisqually in our case study that negative allele counts and frequencies are predicted for one of the minor alleles in the RDA prediction step, but that the GAM step predicts the biologically acceptable frequency of 0.027. Function `AlleleShift::freq.ggplot` (Figure 3) enables a visual inspection of the power of the models to predict allele frequencies for the calibration data.

**Table 3.** Output from function `freq.pred` for the changed climate for four populations and 2 alleles (A and B). This is a subset of the result with all populations and all alleles for the example data sets in *AlleleShift* for *Populus trichocarpa.*

| Population | Puyallup | Tahoe | Skagit | Nisqually |
| --- | --- | --- | --- | --- |
| N | 372 | 24 | 326 | 28 |
| Gene | X01_10838495 |  |  |  |
| Allele.freq | 0.172 | 0.625 | 0.236 | 0.107 |
| A | 64 | 15 | 77 | 3 |
| B | 308 | 9 | 249 | 25 |
| Ap | 144.376 | 23.471 | 75.381 | 15.165 |
| Bp | 227.624 | 0.529 | 250.619 | 12.835 |
| N.e1 | 372 | 24 | 326 | 28 |
| Freq.e1 | 0.388 | 0.978 | 0.231 | 0.542 |
| Freq.e2 | 0.425 | 0.959 | 0.192 | 0.633 |
| LCL | 0.306 | 0.794 | 0.175 | 0.452 |
| UCL | 0.544 | 1.000 | 0.210 | 0.813 |
| increasing | TRUE | TRUE | FALSE | TRUE |
| Gene | X01_16628872 |  |  |  |
| Allele.freq | 0.169 | 0.083 | 0.181 | 0.250 |
| A | 63 | 2 | 59 | 7 |
| B | 309 | 22 | 267 | 21 |
| Ap | 38.751 | 0.509 | 52.668 | -0.038 |
| Bp | 333.249 | 23.491 | 273.332 | 28.038 |
| N.e1 | 372 | 24 | 326 | 28 |
| Freq.e1 | 0.104 | 0.021 | 0.162 | -0.001 |
| Freq.e2 | 0.104 | 0.037 | 0.183 | 0.027 |
| LCL | 0.091 | 0.029 | 0.161 | 0.018 |
| UCL | 0.117 | 0.045 | 0.205 | 0.036 |
| increasing | FALSE | FALSE | TRUE | FALSE |

**Figure 3.**
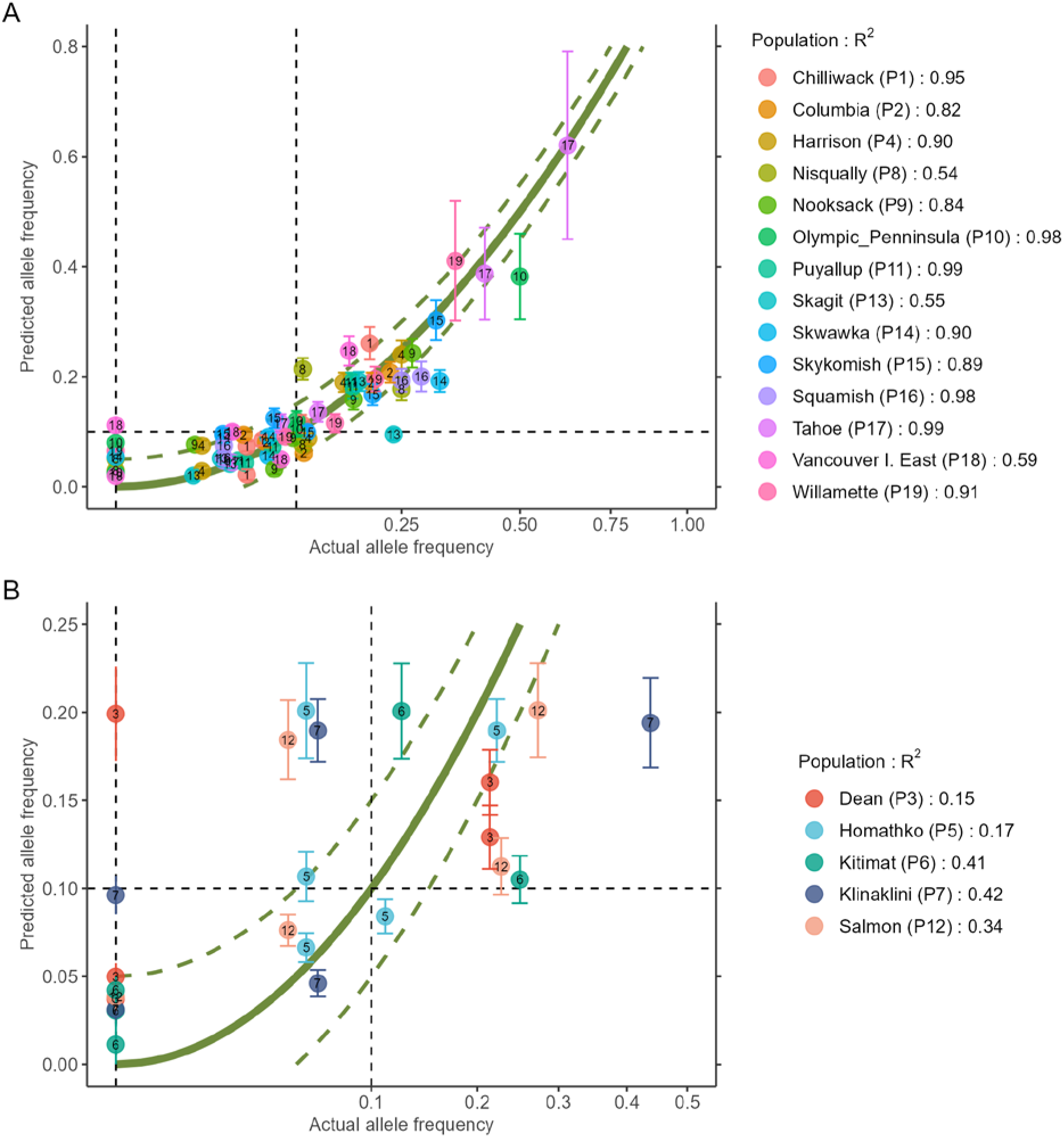
Plot of the actual frequency of the minor allele against the predicted frequency for the calibration data. The ‘olivegreen’ reference lines indicate 1:1 (bold line), 1:1.05 and 1:0.95 (dashed lines) relationships. (A) Data for populations where a linear model explains more than 50% in allele frequencies. (B) Data for populations where a linear model explains less than 50% in allele frequencies.

### Visualizations

*AlleleShift* generates various types of *ggplot2* (Wickham, 2016; version 3.3.2) graphics from the output of `AlleleShift::freq.pred`. These include dot (Figure 4), pie or donut (Figure 5), moon (Figure 6) and waffle (Figure 7) graphics, and smoothed regression surfaces (Figure 8). As an intermediate step to generate various of these graphics, helper functions such as `waffle.baker` for `shift.waffle.ggplot` or `moon.waxer` for `shift.moon.ggplot` prepare data for the main graphing function (Table 2). With default settings, visualizations depict changes in allele frequencies for each allele in different panels, internally via `ggplot2::facet_grid`. Setting argument `mean.change` to `TRUE`, the graphics depict median or mean changes in allele frequencies.

**Figure 4.**
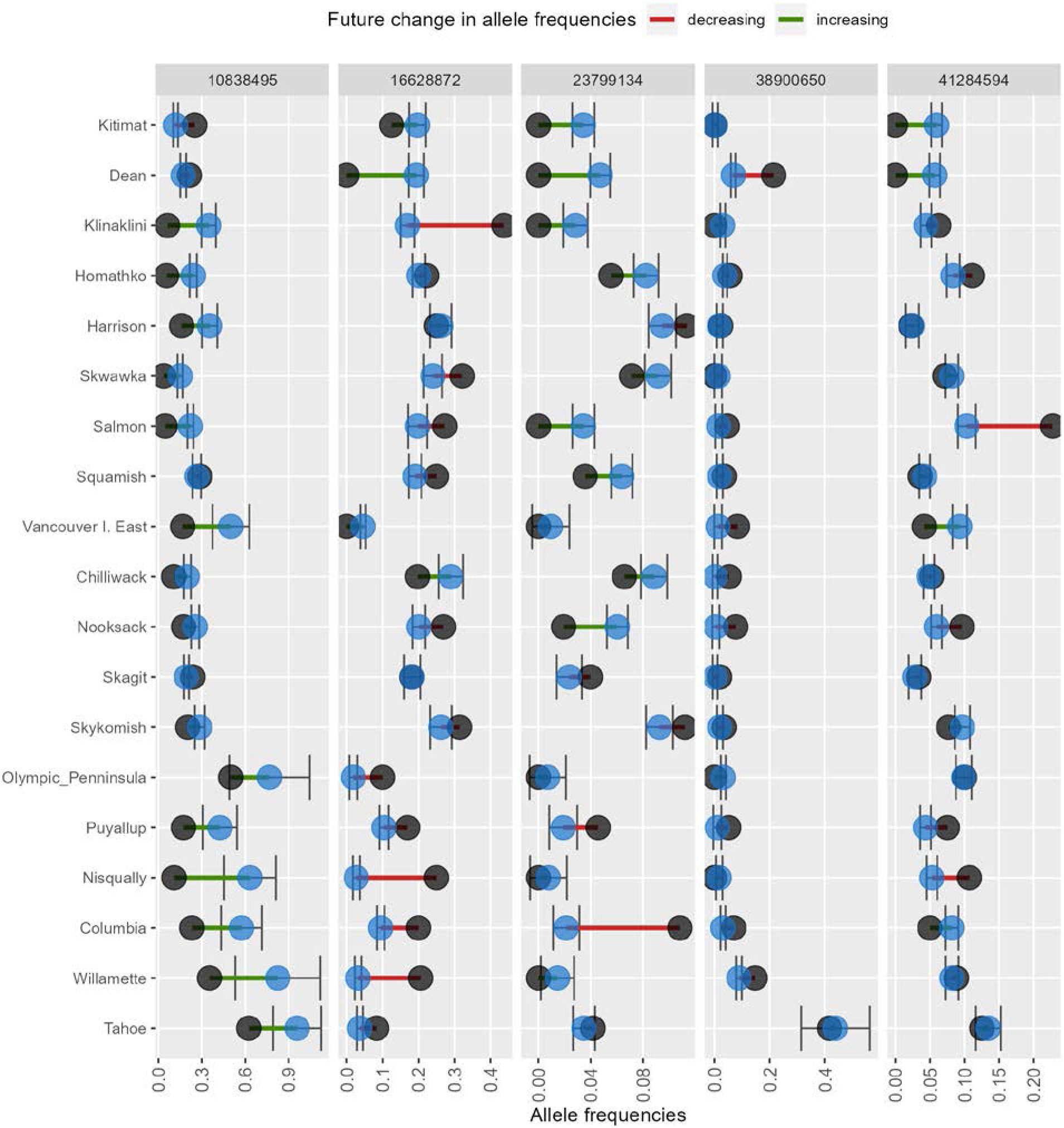
Depiction of changes in minor allele frequencies via. `AlleleShift::shift.dot.ggplot`. Black circles reflect baseline frequencies, blue circles future frequencies and vertical lines the confidence interval

**Figure 5.**
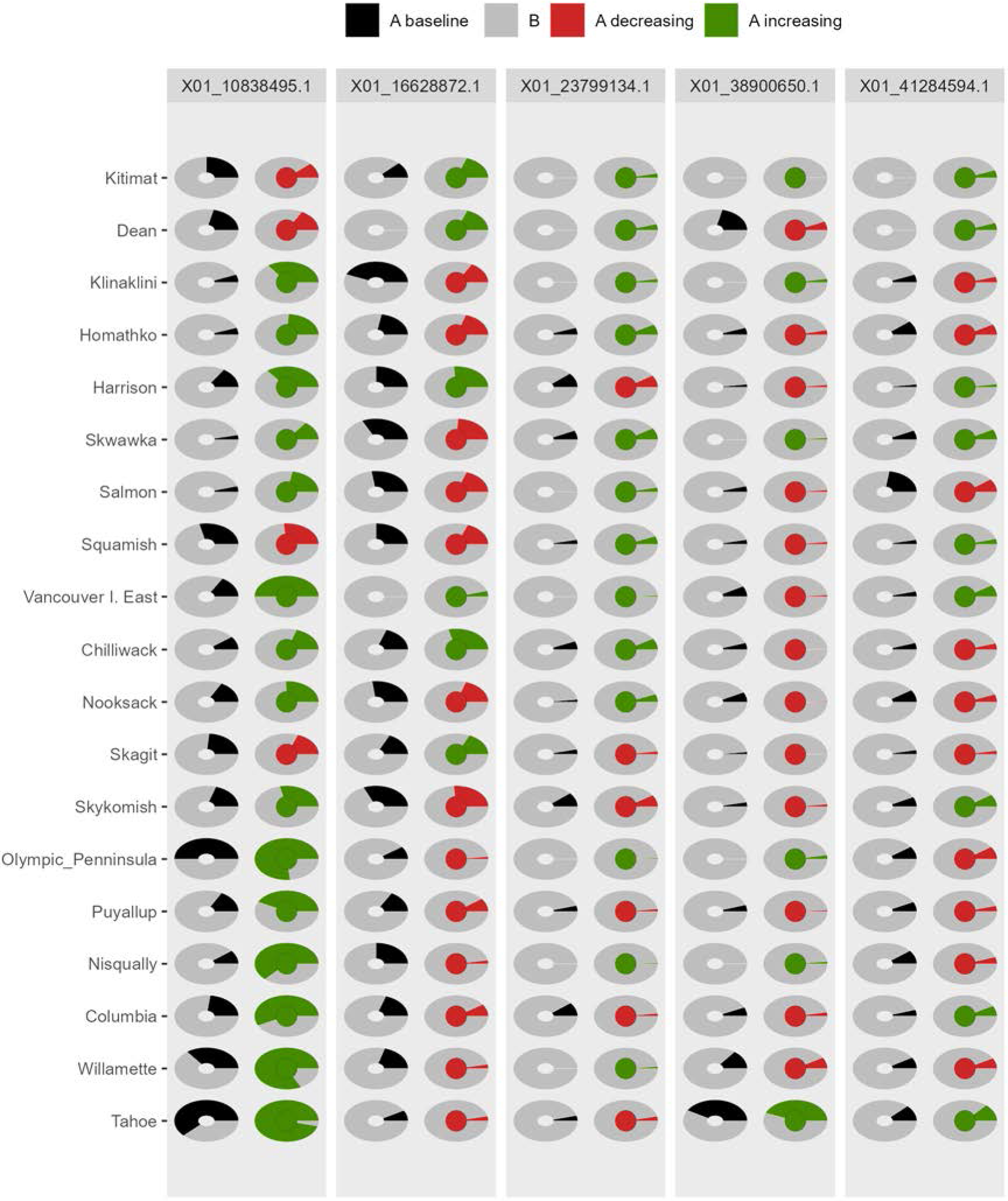
Depiction of changes in allele frequencies via `AlleleShift::shift.pie.ggplot`. Columns on the left reflect baseline frequencies with frequency of the minor allele in black. Columns on the right reflect future frequencies, with colour of the arc and colour of the central circle reflecting frequencies and trends (red = decreasing, green = increasing) of the minor allele.

**Figure 6.**
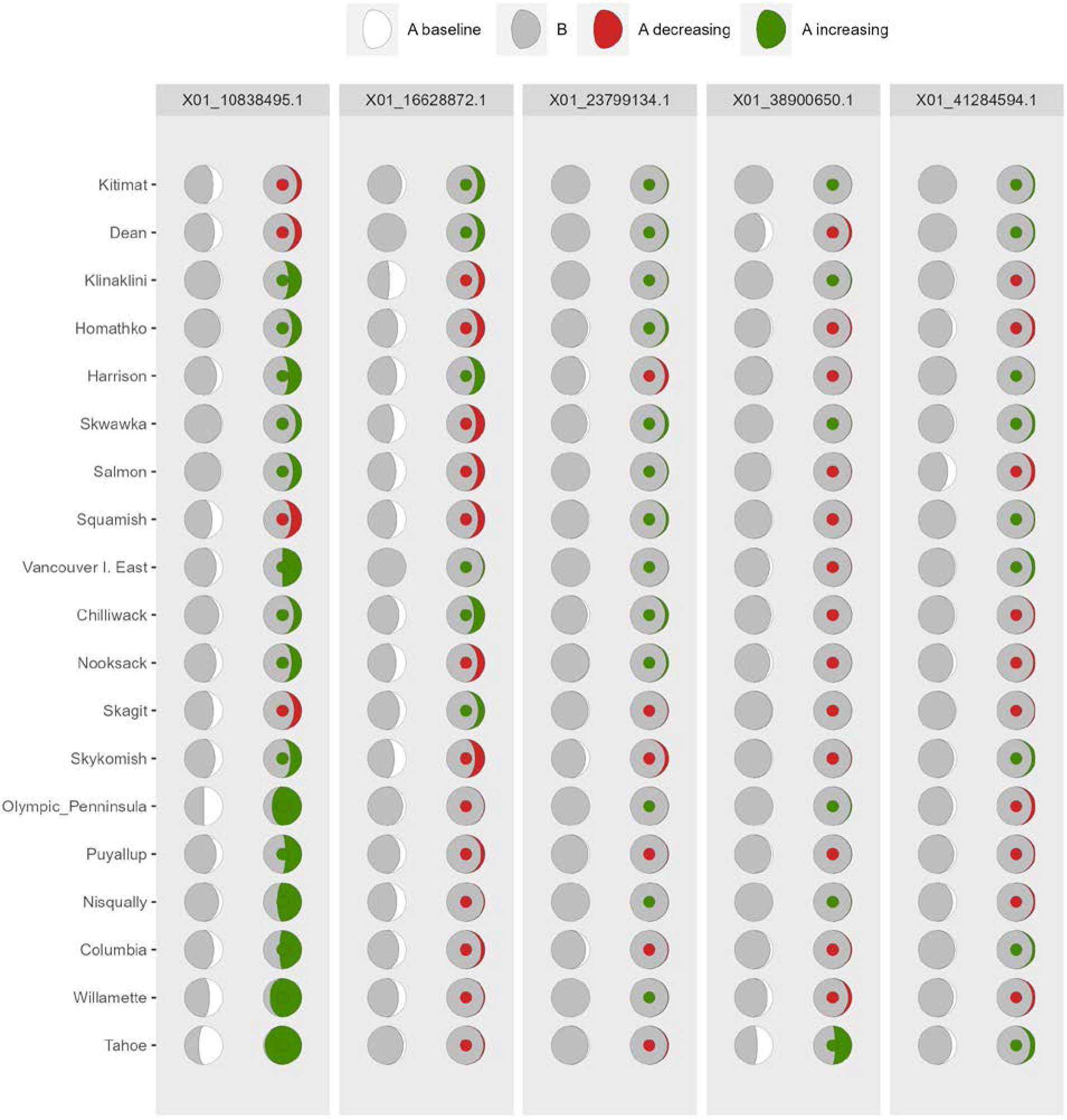
Depiction of changes in allele frequencies via `AlleleShift::shift.moon.ggplot`. Columns on the left reflect baseline frequencies with frequency of the minor allele in white. Columns on the right reflect future frequencies, with colour of the waxing moon and colour of the central circle reflecting frequencies and trends (red = decreasing, green = increasing) of the minor allele.

**Figure 7.**
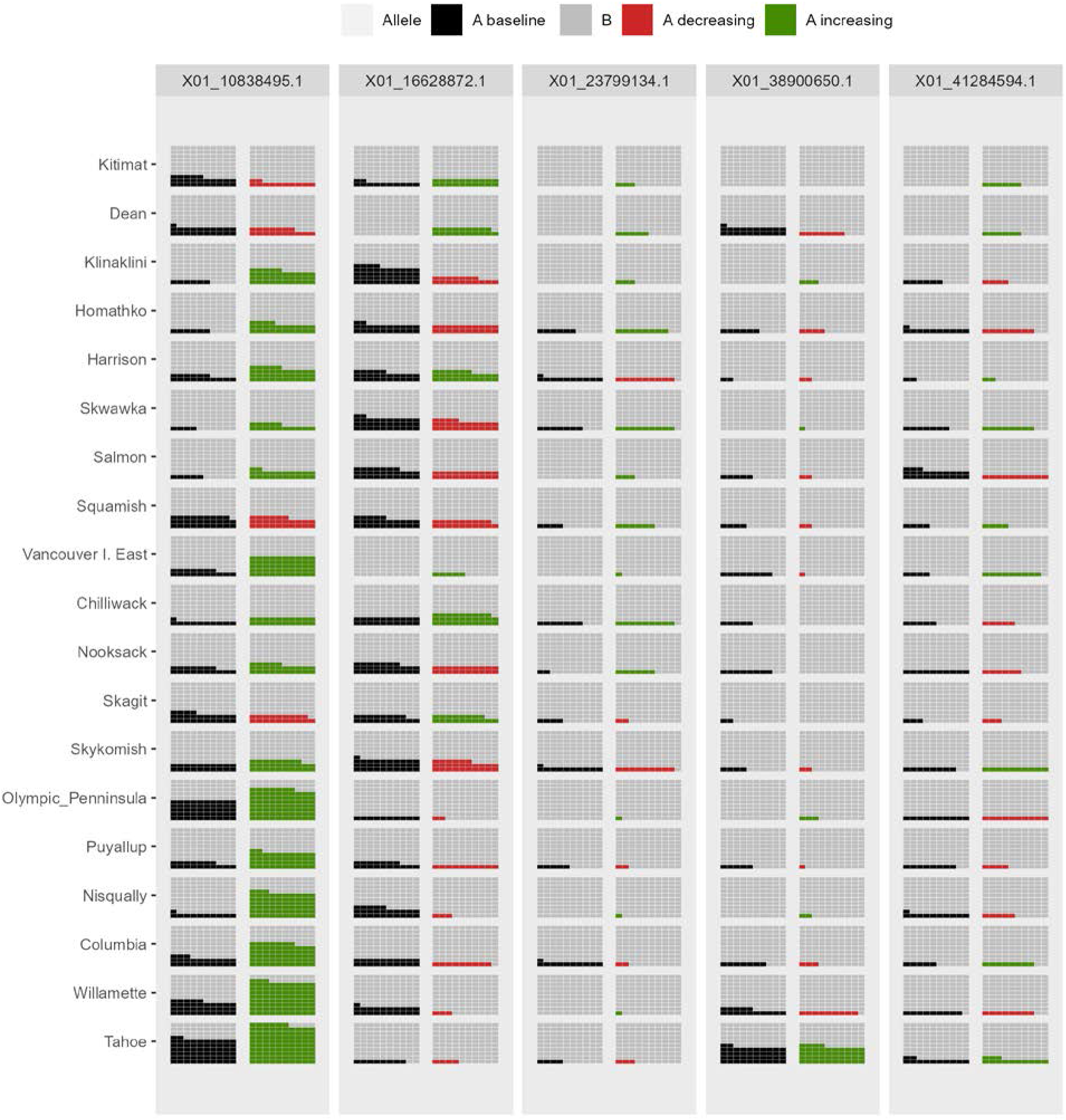
Depiction of changes in allele frequencies via `AlleleShift::shift.waffle.ggplot`. Each ‘waffle’ has 100 ‘cells’. Columns on the left reflect baseline frequencies with frequency of the minor allele in black. Columns on the right reflect future frequencies of the minor allele, with colour indicating trends (red = decreasing, green = increasing)

**Figure 8.**
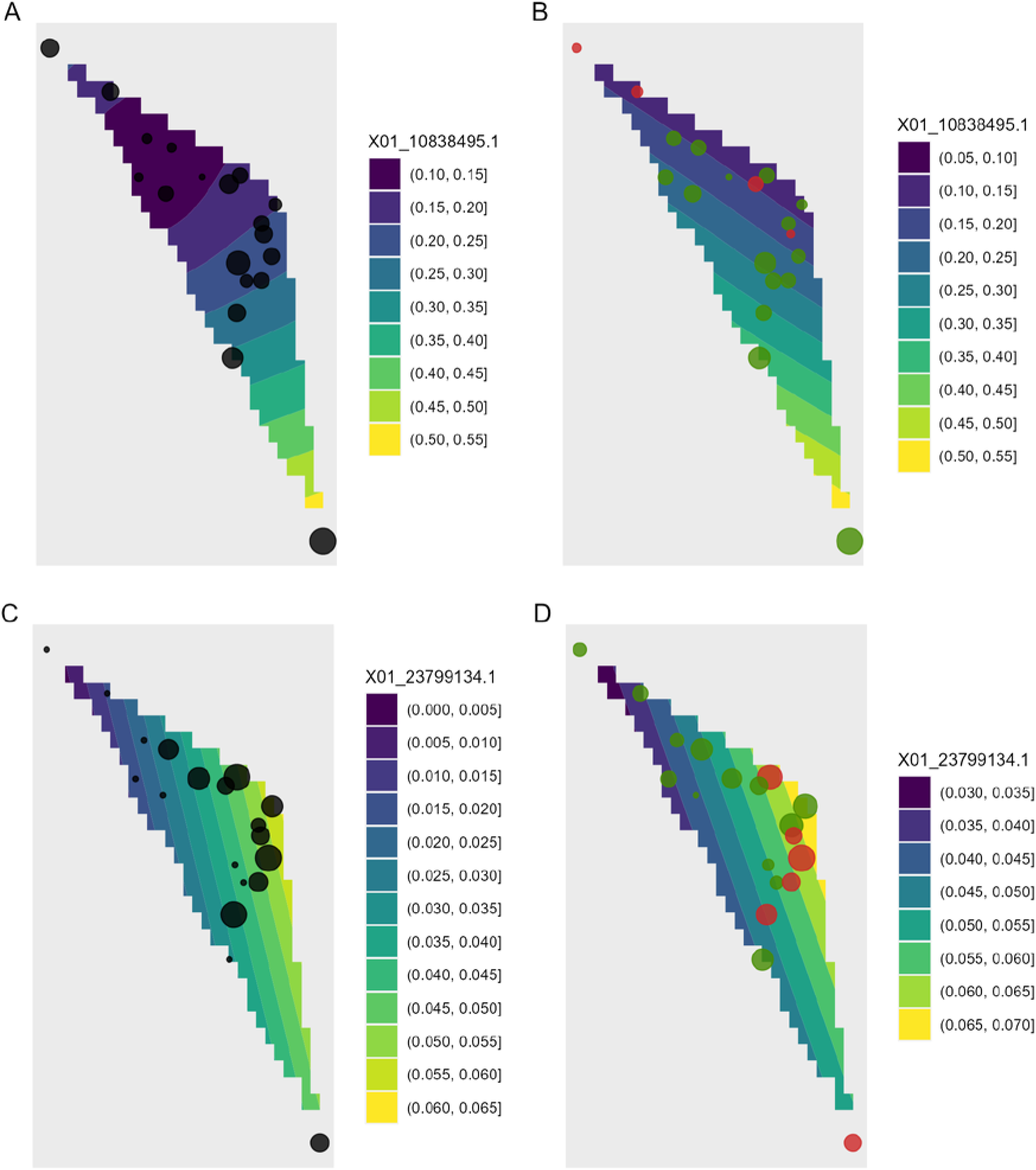
Depiction of changes in allele frequencies in geographical space via `AlleleShift::shift.surf.ggplot`. (A) Frequencies in the baseline climate for the minor allele for locus 10838495. (B) Frequencies in a future climate for the same allele as in (A). Frequencies in the baseline climate for the minor allele for locus 23799134. (D) Frequencies in a future climate for the same allele as in (D). Sizes of circles reflect the frequencies for the populations. Colours for the future frequencies (B, D)_indicate trends (red = decreasing, green = increasing).

Function `shift.surf.ggplot` plots populations in geographical space via their geographical coordinates (longitude and latitude in Figure 8). The function fits a smoothed regression surface for allele frequencies via `vegan::ordisurf`. This output is then further processed internally within the function via `BiodiversityR::ordisurfgrid.long` (an overview of generating *ggplot2* ordination diagrams via *vegan* and *BiodiversityR* is given in Kindt, 2020b; these guidelines can be used to customize ordination graphs as shown in Figures 2 and 8). Various options of fitting smoothed regression surfaces are available through providing additional arguments to `shift.surf.ggplot`, such as the various `mgcv::smooth.terms` options of thin plates, Duchon splines, cubic regression splines, P-splines, Markov Random Fields, Gaussian process smooths, soap film smooths, splines on the sphere and random effects.

As graphics are generated with *ggplot2*, it is relatively easy to generate animated versions of visualizations with *gganimate* (Pedersen & Robinson, 2020; version 1.0.7). Scripts for generating animated versions for dot graphics, pie graphics and smoothed surfaces are provided with the documentation of the respective functions. The supplementary materials show MP4 videos for a time series that interpolates bioclimatic data between baseline and future climate in steps of five years; these also extrapolate data far into the future (up to a million years, see the Results and Discussion for my reason to do this). Other than time series, animated visualizations could also be generated for different global circulation models (GCMs) or scenarios such as various shared socio-economic pathways (SSPs) developed in the context of the sixth assessment report of the Intergovernmental Panel on Climate Change.

## Results and Discussion

*AlleleShift* predicts shifts in allele frequencies via RDA and GAM, an alternative pathway that maintains Euclidean distances among populations and individuals (see Supplementary materials 1). It also avoids making negative frequency predictions as can occur with the CCA approach recently devised by Blumstein et al. (2020). My methodology however faces the same limitations of data requirements as discussed by Blumstein et al. (2020) for their protocol in terms of the initial identification of responsive markers. Key assumptions of the Blumstein et al. (2020) protocol apply also to my approach, including that allelic effects are independent and additive with no epistasis or dominance (see also the discussion on epistasis, structural genomic variation and epigenetics by Stange, Barrett & Hendry, 2020).

I recommend reducing both the explanatory variables and the response variables to subsets of data with a maximum Variance Inflation Factor (VIF) of 20 or less, for alleles to obtain better estimates of changes in their frequencies. For the allele counts as response variables, if VIF would be larger than 20, I would use function `VIF.subset` in an iterative procedure whereby earlier selected subsets of alleles are excluded, to generate different subsets of alleles (but keeping variables for both A and B allele counts in each final subset). For example, with 20 alleles X01A, X01B, X02A, X02B, … , X10A, X10B, the two subsets could be {X01A, X01B, …, X05A, X05B} and {X06A, X06B, … , X10A, X10B}. With the various subsets, shifts in allele frequencies can then be predicted, and finally predictions with all subsets can be combined.

When predictions are made by *AlleleShift* into the future, and especially into novel climatic conditions, it is warranted to consider the transferability of the calibrated models and ideally to provide ‘transferability metrics’ that quantify prediction uncertainty (Yates et al., 2018). For the animated graphics, as an extreme example of the caveats of the methodology, I made projections one million years into the future. These projections followed earlier trends of the 21^st^ century and resulted in allele frequencies becoming fixed at 0 or 1, which is a biologically possible scenario. At the same time, however, these simulations clearly illustrated that the methodology through RDA is correlative as in correlative/phenomenological species distribution models. In fact, the linear extrapolation of climate variables resulted in an environmental data set where the mean annual temperature was above 30,000 degrees, which obviously is a biologically irrelevant scenario. Thus, the method presented here should be used cautiously in novel climates especially as predictions will be available (the model will not crash or give an error warning), and should thus be used even more cautiously where differences between baseline and future climates are large.

As an approach to obtain a better handle on the transferability of *AlleleShift* predictions into future climates, I recommend to also estimate the suitability (and ideally the transferability, for instance via evaluation strips as proposed by Elith et al., 2005) of target species via species distribution models (SDMs). Well-documented methods of utilizing SDMs to predict shifts in species habitat are available in the literature, including recent examples that use the ensemble suitability modelling framework available in *BiodiversityR* (e.g., de Sousa et al., 2019; Fremout et al., 2020; Kindt, 2018; Ranjitkar et al., 2014). For organisms such as trees, correlative SDMs remain the best available method of predicting future species suitability, whereas the limitations of these methods may not be as great as has been suggested (Booth, 2018). What is also attractive about SDMs is that a wider set of presence observations are likely to be available than those populations that have been studied genetically. Presence data are available from open-source databases such as GBIF or the Botanical Information and Ecology Network (Enquist et al., 2016). Further to the collation of a larger set of presence observations, application of SDMs should be straightforward using the same (bio)climatic data sets as applied in *AlleleShift*. The approach of combining the results of *AlleleShift* with SDM is somewhat similar to the method applied by Aguirre-Liguori et al. (2019) to develop species distribution models for alleles. In my proposal, however, the predictions of allele frequencies and SDM are done independently, and ideally with an expanded point presence data set for SDM.

A straightforward and practical expansion of the methodology I have proposed is to tree seed sourcing programmes, possibly for developing schemes of human-assisted geneflow *sensu* Aitken & Whitlock (2013). This is important for ensuring the matching of planting materials to the conditions of planting sites (Cernansky, 2018; Roshetko et al. 2018; Kettle et al. 2020). For specific planting sites and planting times (considering the perennial nature of trees, climate change during the production cycle should be considered) of interest, the prediction methods can readily provide the predicted allele frequencies needed for adaptation. Theoretically, based on the similarity between predicted allele frequencies and those of available source populations, the best matching source can then be selected.

## Conclusions

The R package *AlleleShift* provides a set of functions that allow the prediction of allele frequencies from baseline, future and past (bio)climatic explanatory variables via redundancy analysis (RDA) and generalized additive models (GAM). Various visualizations are provided via *ggplot2* and its extension packages such as *ggforce* and *gganimate*. At the time of submission of this manuscript, no package was available for this set of tools. As with any other methodology that projects into the future, it is important to reflect on transferability to novel climates.

## Supporting information

Supplementary Materials on relationship between Euclidean distances among populations and centroids of individuals

Animated dot graphic for shifts in allele frequencies

Animdated pie graphics for shifts in allele frequencies

Animated smoothed regression surface graphics for shifts in allele frequencies

## Acknowledgements

The author thanks Ian Dawson (CIFOR-ICRAF) for reviewing the article prior to submission. He also thanks additional colleagues from CIFOR-ICRAF and the University of Copenhagen for useful discussions on the applications of this package, including Lars Graudal, Prasad Hendre, Ramni Jamnadass and Jens-Peter B. Lillesø. The CGIAR Research Program on Forests, Trees, and Agroforestry (supported by the CGIAR Trust Fund) and the Provision of Adequate Tree Seed Portfolios project (supported by Norway’s International Climate and Forest Initiative through the Royal Norwegian Embassy in Ethiopia) supported the author’s time on this project.

## References

Aguirre-Liguori, J.A., Ramírez-Barahona, S., Tiffin, P. and Eguiarte, L.E., 2019. Climate change is predicted to disrupt patterns of local adaptation in wild and cultivated maize. Proceedings of the Royal Society B, 286(1906), p.20190486. https://doi.org/10.1098/rspb.2019.0486

Aitken, S.N., Whitlock, M.C. (2013). Assisted gene flow to facilitate local adaptation to climate change. Annual review of ecology, evolution, and systematics, 44. https://doi.org/10.1146/annurev-ecolsys-110512-135747

Anderson, J.T. and Song, B.-H. (2020). Plant adaptation to climate change—Where are we?. J. Syst. Evol., 58: 533–545. https://doi.org/10.1111/jse.12649

Blumstein et al. 2020. Protocol for Projecting Allele Frequency Change under Future Climate Change at Adaptive-Associated Loci. https://doi.org/10.1016/j.xpro.2020.100061

Booth, T. H. (2018). Species distribution modelling tools and databases to assist managing forests under climate change. Forest Ecology and Management 430: 196–203. URL https://doi.org/10.1016/j.foreco.2018.08.019.

Booth, T.H. (2016). Estimating potential range and hence climatic adaptability in selected tree species. Forest Ecology and Management, 366, pp.175–183. https://doi.org/10.1016/j.foreco.2016.02.009

Bramson, M. (2019). gggibbous: Moon Charts, a Pie Chart Alternative. R package version 0.1.0. https://CRAN.R-project.org/package=gggibbous

Brown, J., Hill, D., Dolan, A. et al. (2018). PaleoClim, high spatial resolution paleoclimate surfaces for global land areas. Sci Data 5, 180254. https://doi.org/10.1038/sdata.2018.254

Capblancq, T, Morin, X, Gueguen, M, Renaud, J, Lobreaux, S, Bazin, E. Climate–associated genetic variation in Fagus sylvatica and potential responses to climate change in the French Alps. J Evol Biol. 2020; 33: 783–796. https://doi.org/10.1111/jeb.13610

Cernansky, R. (2018). How to plant a trillion trees. Nature 560: 542–544. https://doi.org/10.1038/d41586-018-06031-x

de Sousa, K., van Zonneveld, M., Holmgren, M. et al. (2019). The future of coffee and cocoa agroforestry in a warmer Mesoamerica. Sci Rep 9, 8828. https://doi.org/10.1038/s41598-019-45491-7

Elith, J., Ferrier, S., Huettmann, F. and Leathwick, J., 2005. The evaluation strip: a new and robust method for plotting predicted responses from species distribution models. Ecological modelling, 186(3), pp.280–289. https://doi.org/10.1016/j.ecolmodel.2004.12.007

Enquist, B.J., Condit, R., Peet, R.K., Schildhauer, M. and Thiers, B.M., 2016. Cyberinfrastructure for an integrated botanical information network to investigate the ecological impacts of global climate change on plant biodiversity (No. e2615v2). PeerJ Preprints. https://doi.org/10.7287/peerj.preprints.2615v2

Excoffier, L., Smouse, P.E. and Quattro, J.M. (1992). Analysis of molecular variance inferred from metric distances among DNA haplotypes: application to human mitochondrial DNA restriction data. Genetics, 131, 479–491.

Fick, S.E., Hijmans, R.J. (2017). WorldClim 2: new 1km spatial resolution climate surfaces for global land areas. International Journal of Climatology 37 (12): 4302–4315. https://www.worldclim.org

Fox, J., Monette, G., 1992. Generalized collinearity diagnostics. J. Am. Stat. Assoc. 87, 178e183. https://doi.org/10.1080/01621459.1992.10475190.

Fremout, T, Thomas, E, Gaisberger, H, et al. (2020). Mapping tree species vulnerability to multiple threats as a guide to restoration and conservation of tropical dry forests. Glob Change Biol.; 26: 3552–3568. https://doi.org/10.1111/gcb.15028

Günther, T. and Coop, G., 2013. Robust identification of local adaptation from allele frequencies. Genetics, 195(1), pp.205–220. https://dx.doi.org/10.1534%2Fgenetics.113.152462

Kamvar ZN, Tabima JF, Grünwald NJ. (2014) Poppr: an R package for genetic analysis of populations with clonal, partially clonal, and/or sexual reproduction. PeerJ 2:e281. https://doi.org/10.7717/peerj.281

Kettle, C.J., Atkinson, R., Boshier, D., Ducci, F., Dawson, I., Ekué, M., Elias, M., Graudal, L., Jalonen, R., Koskela, J. and Monteverdi, M.C., 2020. Priorities, challenges and opportunities for supplying tree genetic resources. Restoring the Earth-The next decade: Unasylva No. 252-Vol. 71 2020/1, 252(1), p.51. http://www.fao.org/3/cb1600en/CB1600EN.pdf

Kindt, R. (2018). Ensemble species distribution modelling with transformed suitability values. Environmental Modelling & Software 100: 136–145. https://doi.org/10.1016/j.envsoft.2017.11.009

Kindt, R. (2020a). Ordination graphs with vegan, BiodiversityR and ggplot2. https://rpubs.com/Roeland-KINDT

Kindt, R. (2020b). Analysis of Molecular Variance (AMOVA) with vegan and BiodiversityR, including a graphical method to identify potential migrants. https://rpubs.com/Roeland-KINDT

Kindt, R., Coe, R. (2005). Tree Diversity Analysis. A manual and software for common statistical methods for ecological and biodiversity studies. https://CRAN.R-project.org/package=BiodiversityR

Lasky, J.R., Forester, B.R. and Reimherr, M., 2018. Coherent synthesis of genomic associations with phenotypes and home environments. Molecular Ecology Resources, 18(1), pp.91–106. https://doi.org/10.1111/1755-0998.12714

Legendre, P., Legendre, L., 2012. Numerical ecology. Elsevier.

Luikart G., Kardos M., Hand B.K., Rajora O.P., Aitken S.N., Hohenlohe P.A. (2018) Population Genomics: Advancing Understanding of Nature. In: Rajora O. (eds) Population Genomics. Population Genomics. Springer, Cham. https://doi.org/10.1007/13836_2018_60

Meirmans, P., Liu, S. (2018). Analysis of Molecular Variance (AMOVA) for Autopolyploids Front. Ecol. Evol., 23. https://doi.org/10.3389/fevo.2018.00066

Meybeck, A., Gitz, V., Wolf, J. and Wong, T. 2020. Addressing forestry and agroforestry in National Adaptation Plans – Supplementary guidelines. Place of publication, Bogor/Rome. FAO and FTA. https://doi.org/10.4060/cb1203en

Michalakis, Y., Excoffier, L. (1996). A generic estimation of population subdivision using distances between alleles with special reference for microsatellite loci. Genetics 142, 1061–1064.

Nelson, J.T., Motamayor, J.C. and Cornejo, O.E., 2020. Environment and pathogens shape local and regional adaptations to climate change in the chocolate tree, *Theobroma cacao* L. Molecular Ecology. https://doi.org/10.1111/mec.15754

Peakall R and Smouse PE. 2012. GenAlEx 6.5: genetic analysis in Excel. Population genetic software for teaching and research - an update. https://doi.org/10.1093/bioinformatics/bts460.

Pedersen, T.L. (2020). ggforce: Accelerating ‘ggplot2’. R package version 0.3.2. https://CRAN.R-project.org/package=ggforce

Pedersen, T.L., Robinson, D. (2020). gganimate: A Grammar of Animated Graphics. R package version 1.0.7. https://CRAN.R-project.org/package=gganimate

Ranjitkar, S., Xu, J., Shrestha, K.K. and Kindt, R. (2014). Ensemble forecast of climate suitability for the Trans-Himalayan Nyctaginaceae species. Ecological Modelling, 282: 8–24. https://doi.org/10.1016/j.ecolmodel.2014.03.003

Razgour, O., Forester, B., Taggart, J.B., Bekaert, M., Juste, J., Ibáñez, C., Puechmaille, S.J., Novella-Fernandez, R., Alberdi, A. and Manel, S. (2019). Considering adaptive genetic variation in climate change vulnerability assessment reduces species range loss projections. Proceedings of the National Academy of Sciences, 116(21), 10418–10423. https://doi.org/10.1073/pnas.1820663116

Rengefors, K., Gollnisch, R., Sassenhagen, I., Härnström Aloisi, K., Svensson, M., Lebret, K., Čertnerová, D., Cresko, W.A., Bassham, S. and Ahrén, D. (2021). Genome–wide SNP markers reveal population structure and dispersal direction of an expanding nuisance algal bloom species. Molecular Ecology. https://doi.org/10.1111/mec.15787

Ripple W.J., Christopher Wolf, Thomas M Newsome, Phoebe Barnard, William R Moomaw, et al. (2020). World Scientists’ Warning of a Climate Emergency, BioScience 70: 8–12, https://doi.org/10.1093/biosci/biz088

Roshetko, J.M., Dawson, I.K., Urquiola, J., Lasco, R.D., Leimona, B., Weber, J.C., Bozzano, M., Lillesø, J.P.B., Graudal, L. and Jamnadass, R. (2018). To what extent are genetic resources considered in environmental service provision? A case study based on trees and carbon sequestration. Climate and Development, 10(8), pp.755–768. https://doi.org/10.1080/17565529.2017.1334620

Stange, M., Barrett, R.D. and Hendry, A.P. (2020). The importance of genomic variation for biodiversity, ecosystems and people. Nature Reviews Genetics, pp.1–17.

Stanturf, J.A, Kant, P., Lillesø, J.-P.B., Mansourian, S., Kleine, M., Lars Graudal, L., Palle Madsen, P. (2015). Forest Landscape Restoration as a Key Component of Climate Change Mitigation and Adaptation. IUFRO World Series Volume 34. Vienna 72 p.

Temunović, M., Garnier–Géré, P., Morić, M., Franjić, J., Ivanković, M., Bogdan, S. and Hampe, A. (2020). Candidate gene SNP variation in floodplain populations of pedunculate oak (*Quercus robur* L.) near the species’ southern range margin: Weak differentiation yet distinct associations with water availability. Molecular Ecology, 29(13), pp.2359–2378. https://doi.org/10.1111/mec.15492

Title P.O., Bemmels J.B. (2018). ENVIREM: an expanded set of bioclimatic and topographic variables increases flexibility and improves performance of ecological niche modeling. Ecography. 41:291–307. https://envirem.github.io/

Waldvogel, A.-M., Feldmeyer, B., Rolshausen, G., Exposito-Alonso, M., Rellstab, C., Kofler, R., Mock, T., Schmid, K., Schmitt, I., Bataillon, T., Savolainen, O., Bergland, A., Flatt, T., Guillaume, F. and Pfenninger, M. (2020), Evolutionary genomics can improve prediction of species’ responses to climate change. Evolution Letters, 4: 4–18. https://doi.org/10.1002/evl3.154

Wickham, H. (2016). ggplot2: Elegant Graphics for Data Analysis. Springer-Verlag New York, 2016. https://ggplot2.tidyverse.org

Wood, S.N. (2004) Stable and Efficient Multiple Smoothing Parameter Estimation for Generalized Additive Models, Journal of the American Statistical Association, 99: 467, 673–686. https://doi.org/10.1198/016214504000000980

Yates, K.L., Phil J. Bouchet, P.J., Caley, M.J., Mengersen, K., Randin, C.F., Parnell, S., Fielding, A.H., Bamford, A.J., Ban, S., et al. (2018). Outstanding Challenges in the Transferability of Ecological Models. Trends in Ecology & Evolution 33: 790–802. URL https://doi.org/10.1016/j.tree.2018.08.001.

