## Supplementary Materials on relationship between Euclidean distances among populations and centroids of individuals for "*AlleleShift*: An R package to predict and visualize population-level changes in allele frequencies in response to climate change"

### Population and individual genetic distance

Roeland Kindt

10/01/2021

#### 2. Packages needed

```
library(BiodiversityR) # also loads vegan
library(poppr) # also loads adegenet
library(AlleleShift)
```

#### 3. Objective

Here I show how Euclidean distances among populations directly correspond to Euclidean distances among the Population centroids that are obtained from the Principal Component Analysis (PCA) scores of individuals.

#### 4. Load example data set

The **AlleleShift** package contains a data set with allele counts for 902 individuals that belong to 19 populations.

```
data(Poptri.genind)
poppr::poppr(Poptri.genind)

##                  Pop   N MLG eMLG    SE     H    G lambda   E.5  Hexp       Ia
## 1         Chilliwack  38  14 5.67 1.244 1.972 3.97  0.748 0.480 0.168  0.29948
## 2           Columbia 180  31 6.69 1.276 2.634 8.17  0.878 0.555 0.219  0.03214
## 3               Dean   7   4 4.00 0.000 1.352 3.77  0.735 0.967 0.145 -0.16667
## 4           Harrison  22  10 5.87 1.093 1.855 4.10  0.756 0.575 0.191  0.06387
## 5           Homathko   9   7 7.00 0.000 1.831 5.40  0.815 0.840 0.182 -0.17703
## 6            Kitimat   8   4 4.00 0.000 1.321 3.56  0.719 0.930 0.127 -0.13245
## 7         Klinaklini   8   5 5.00 0.000 1.494 4.00  0.750 0.868 0.155 -0.41248
## 8          Nisqually  14   7 5.51 0.830 1.567 3.38  0.704 0.627 0.157  0.00845
## 9           Nooksack  26  11 6.44 1.072 2.124 6.76  0.852 0.782 0.211  0.05709
## 10 Olympic_Penninsula   5   3 3.00 0.000 0.950 2.27  0.560 0.802 0.191 -0.25000
## 11          Puyallup 186  29 6.03 1.303 2.396 5.81  0.828 0.482 0.178 -0.02222
## 12            Salmon  11   7 6.64 0.481 1.846 5.76  0.826 0.892 0.193 -0.19978
## 13            Skagit 163  20 5.41 1.268 2.102 4.72  0.788 0.518 0.168 -0.01194
## 14           Skwawka  14   6 5.29 0.656 1.631 4.45  0.776 0.841 0.160 -0.04804
## 15          Skykomish 156  31 6.97 1.249 2.730 9.26  0.892 0.576 0.233  0.07931
## 16           Squamish  14  10 7.86 0.802 2.206 8.17  0.878 0.887 0.205 -0.28884
## 17             Tahoe  12   9 7.82 0.649 2.095 7.20  0.861 0.870 0.293 -0.11481
## 18  Vancouver I. East  12   4 3.50 0.584 0.837 1.71  0.417 0.546 0.107  0.48361
## 19         Willamette  17  10 6.65 1.016 2.007 5.45  0.817 0.691 0.246  0.11902
## 20             Total 902  59 6.38 1.323 2.663 6.93  0.856 0.445 0.200  0.03621
##       rbarD          File
## 1   0.08166 Poptri.genind
## 2   0.00883 Poptri.genind
## 3  -0.16667 Poptri.genind
## 4   0.02067 Poptri.genind
## 5  -0.04770 Poptri.genind
## 6  -0.14222 Poptri.genind
## 7  -0.21444 Poptri.genind
## 8   0.00445 Poptri.genind
## 9   0.01646 Poptri.genind
## 10 -0.25000 Poptri.genind
## 11 -0.00606 Poptri.genind
## 12 -0.07663 Poptri.genind
## 13 -0.00373 Poptri.genind
```

```
## 14 -0.01849 Poptri.genind
## 15  0.02249 Poptri.genind
## 16 -0.08273 Poptri.genind
## 17 -0.03060 Poptri.genind
## 18  0.27081 Poptri.genind
## 19  0.04139 Poptri.genind
## 20  0.00998 Poptri.genind
```

### 5. Distances among populations directly calculated from a genpop object

#### 5.1. Create the genpop object

```
Poptri.genpop <- genind2genpop(Poptri.genind)

##
##  Converting data from a genind to a genpop object...
##
## ...done.

Poptri.genpop

## /// GENPOP OBJECT /////////
##
##  // 19 populations; 5 loci; 10 alleles; size: 8 Kb
##
##  // Basic content
##    @tab:  19 x 10 matrix of allele counts
##    @loc.n.all: number of alleles per locus (range: 2-2)
##    @loc.fac: locus factor for the 10 columns of @tab
##    @all.names: list of allele names for each locus
##    @ploidy: ploidy of each individual  (range: 2-2)
##    @type:  codom
##    @call: genind2genpop(x = Poptri.genind)
##
##  // Optional content
##    - empty -

adegenet::makefreq(Poptri.genpop)

##
##  Finding allelic frequencies from a genpop object...
##
## ...done.

##                      X01_10838495.1 X01_10838495.2 X01_16628872.1 X01_16628872.2
## Chilliwack               0.10526316      0.8947368     0.19736842      0.8026316
## Columbia                 0.23055556      0.7694444     0.20000000      0.8000000
## Dean                     0.21428571      0.7857143     0.00000000      1.0000000
## Harrison                 0.15909091      0.8409091     0.25000000      0.7500000
## Homathko                 0.05555556      0.9444444     0.22222222      0.7777778
## Kitimat                  0.25000000      0.7500000     0.12500000      0.8750000
## Klinaklini               0.06250000      0.9375000     0.43750000      0.5625000
## Nisqually                0.10714286      0.8928571     0.25000000      0.7500000
## Nooksack                 0.17307692      0.8269231     0.26923077      0.7307692
## Olympic_Penninsula       0.50000000      0.5000000     0.10000000      0.9000000
## Puyallup                 0.17204301      0.8279570     0.16935484      0.8306452
## Salmon                   0.04545455      0.9545455     0.27272727      0.7272727
## Skagit                   0.23619632      0.7638037     0.18098160      0.8190184
```

```
## Skwawka                   0.03571429      0.9642857     0.32142857      0.6785714
## Skykomish                 0.20192308      0.7980769     0.31410256      0.6858974
## Squamish                  0.28571429      0.7142857     0.25000000      0.7500000
## Tahoe                     0.62500000      0.3750000     0.08333333      0.9166667
## Vancouver I. East         0.16666667      0.8333333     0.00000000      1.0000000
## Willamette                0.35294118      0.6470588     0.20588235      0.7941176
##                       X01_23799134.1 X01_23799134.2 X01_38900650.1 X01_38900650.2
## Chilliwack                0.06578947      0.9342105     0.05263158      0.9473684
## Columbia                  0.10833333      0.8916667     0.06944444      0.9305556
## Dean                      0.00000000      1.0000000     0.21428571      0.7857143
## Harrison                  0.11363636      0.8863636     0.02272727      0.9772727
## Homathko                  0.05555556      0.9444444     0.05555556      0.9444444
## Kitimat                   0.00000000      1.0000000     0.00000000      1.0000000
## Klinaklini                0.00000000      1.0000000     0.00000000      1.0000000
## Nisqually                 0.00000000      1.0000000     0.00000000      1.0000000
## Nooksack                  0.01923077      0.9807692     0.07692308      0.9230769
## Olympic_Penninsula        0.00000000      1.0000000     0.00000000      1.0000000
## Puyallup                  0.04569892      0.9543011     0.05107527      0.9489247
## Salmon                    0.00000000      1.0000000     0.04545455      0.9545455
## Skagit                    0.03987730      0.9601227     0.01840491      0.9815951
## Skwawka                   0.07142857      0.9285714     0.00000000      1.0000000
## Skykomish                 0.11217949      0.8878205     0.03525641      0.9647436
## Squamish                  0.03571429      0.9642857     0.03571429      0.9642857
## Tahoe                     0.04166667      0.9583333     0.41666667      0.5833333
## Vancouver I. East         0.00000000      1.0000000     0.08333333      0.9166667
## Willamette                0.00000000      1.0000000     0.14705882      0.8529412
##                       X01_41284594.1 X01_41284594.2
## Chilliwack                0.05263158      0.9473684
## Columbia                  0.05000000      0.9500000
## Dean                      0.00000000      1.0000000
## Harrison                  0.02272727      0.9772727
## Homathko                  0.11111111      0.8888889
## Kitimat                   0.00000000      1.0000000
## Klinaklini                0.06250000      0.9375000
## Nisqually                 0.10714286      0.8928571
## Nooksack                  0.09615385      0.9038462
## Olympic_Penninsula        0.10000000      0.9000000
## Puyallup                  0.07526882      0.9247312
## Salmon                    0.22727273      0.7727273
## Skagit                    0.03374233      0.9662577
## Skwawka                   0.07142857      0.9285714
## Skykomish                 0.07692308      0.9230769
## Squamish                  0.03571429      0.9642857
## Tahoe                     0.12500000      0.8750000
## Vancouver I. East         0.04166667      0.9583333
## Willamette                0.08823529      0.9117647
```

#### 5.2. Standardize the data so that each population has the same number of individuals

Data needs to be standardized so that each population has the same number of individuals. If not, the same problem emerges where abundance differences influence the results as in the analysis of differences in species composition through the Euclidean distances that is explained for Figure 8.2 for the **Tree Diversity Analysis** community ecology and biodiversity analysis manual (Kindt and Coe 2005).

Data would also be standardized through argument **scale=TRUE** in **vegan::rda**. But here I use the same standardization pathway that I use in **AlleleShift::count.model**.

The below script standardizes the data so that each population has 186 individuals and 372 alleles. The number of 186 individuals corresponds to the number of individuals in the Puyallup population.

```
gen.comm <- data.frame(Poptri.genpop@tab)

scale2 <- max(rowSums(gen.comm))
scale2/10 # 5 loci with 2 alleles each

## [1] 186

row.ratios <- rowSums(gen.comm) / max(rowSums(gen.comm))
gen.comm.b <- gen.comm
for (i in 1:nrow(gen.comm)) {gen.comm.b[i,] <- gen.comm[i, ] / row.ratios[i]}

rowSums(gen.comm.b[, c(1:2)])

##         Chilliwack          Columbia              Dean          Harrison
##                372               372               372               372
##           Homathko           Kitimat        Klinaklini         Nisqually
##                372               372               372               372
##           Nooksack Olympic_Penninsula         Puyallup            Salmon
##                372               372               372               372
##             Skagit           Skwawka         Skykomish          Squamish
##                372               372               372               372
##              Tahoe  Vancouver I. East        Willamette
##                372               372               372

rowSums(gen.comm.b[, c(3:4)])

##         Chilliwack          Columbia              Dean          Harrison
##                372               372               372               372
##           Homathko           Kitimat        Klinaklini         Nisqually
##                372               372               372               372
##           Nooksack Olympic_Penninsula         Puyallup            Salmon
##                372               372               372               372
##             Skagit           Skwawka         Skykomish          Squamish
##                372               372               372               372
##              Tahoe  Vancouver I. East        Willamette
##                372               372               372

rowSums(gen.comm.b[, c(5:6)])

##         Chilliwack          Columbia              Dean          Harrison
##                372               372               372               372
##           Homathko           Kitimat        Klinaklini         Nisqually
##                372               372               372               372
##           Nooksack Olympic_Penninsula         Puyallup            Salmon
##                372               372               372               372
##             Skagit           Skwawka         Skykomish          Squamish
##                372               372               372               372
##              Tahoe  Vancouver I. East        Willamette
##                372               372               372

rowSums(gen.comm.b[, c(7:8)])
```

```
##         Chilliwack           Columbia               Dean           Harrison 
##                372                372                372                372 
##           Homathko            Kitimat         Klinaklini          Nisqually 
##                372                372                372                372 
##           Nooksack Olympic_Penninsula           Puyallup             Salmon 
##                372                372                372                372 
##             Skagit            Skwawka          Skykomish           Squamish 
##                372                372                372                372 
##              Tahoe  Vancouver I. East         Willamette 
##                372                372                372
```

```
rowSums(gen.comm.b[, c(9:10)])
```

```
##         Chilliwack           Columbia               Dean           Harrison 
##                372                372                372                372 
##           Homathko            Kitimat         Klinaklini          Nisqually 
##                372                372                372                372 
##           Nooksack Olympic_Penninsula           Puyallup             Salmon 
##                372                372                372                372 
##             Skagit            Skwawka          Skykomish           Squamish 
##                372                372                372                372 
##              Tahoe  Vancouver I. East         Willamette 
##                372                372                372
```

#### 5.3. Distances among populations

Now the Euclidean distances among the populations can be calculated.

```
baseline.dist.scaled <- vegdist(gen.comm.b, method="euclidean")
as.matrix(baseline.dist.scaled)[1:7, 1:7]
```

```
##            Chilliwack  Columbia     Dean  Harrison  Homathko   Kitimat
## Chilliwack    0.00000  70.19806 152.5387  51.93464  42.80974  99.89341
## Columbia     70.19806   0.00000 144.5351  54.06585 102.32609  83.28436
## Dean        152.53870 144.53511   0.0000 178.92454 178.56414 131.85695
## Harrison     51.93464  54.06585 178.9245   0.00000  81.08227 102.33144
## Homathko     42.80974 102.32609 178.5641  81.08227   0.00000 134.92817
## Kitimat      99.89341  83.28436 131.8569 102.33144 134.92817   0.00000
## Klinaklini  135.85651 167.49406 270.4467 128.32401 123.29836 194.52346
##            Klinaklini
## Chilliwack   135.8565
## Columbia     167.4941
## Dean         270.4467
## Harrison     128.3240
## Homathko     123.2984
## Kitimat      194.5235
## Klinaklini     0.0000
```

#### 5.4. Distances among populations without standardization

In case the data is not standardized, then the calculations show that population Chilliwack is closer to Dean (distance: 126.8) than to Columbia (distance: 557.2). The opposite is the case in the calculations with the standardized distances directly above. Compare also with the results of section with the Rogers' and Nei distances of section 8.1, where distance Chilliwack-Columbia < Chilliwack-Dean as with the standardized data.

```
baseline.dist2 <- vegdist(Poptri.genpop@tab, method="euclidean")
as.matrix(baseline.dist2)[1:7, 1:7]
```

```
##            Chilliwack Columbia       Dean  Harrison  Homathko    Kitimat
## Chilliwack    0.00000 557.1768 126.822711  66.97761 118.68446 122.000000
## Columbia    557.17681   0.0000 683.047583 623.16290 675.36361 678.213831
## Dean        126.82271 683.0476   0.000000  60.77829  10.67708   6.928203
## Harrison     66.97761 623.1629  60.778286   0.00000  52.45951  55.587768
## Homathko    118.68446 675.3636  10.677078  52.45951   0.00000   6.782330
## Kitimat     122.00000 678.2138   6.928203  55.58777   6.78233   0.000000
## Klinaklini  122.82508 679.3997  11.575837  56.39149   6.63325   8.366600
##            Klinaklini
## Chilliwack  122.82508
## Columbia    679.39974
## Dean         11.57584
## Harrison     56.39149
## Homathko      6.63325
## Kitimat       8.36660
## Klinaklini    0.00000
```

### 6. Distances among centroids of individuals

#### 6.1. Euclidean distances among individuals

```
ind.distA <- as.matrix(vegdist(Poptri.genind@tab,
                  method="euclidean"))[1:7, 1:7]

ind.distA
```

```
##            BESC-52  BESC-79 BESC-246 CA-05-06 BESC-460 GW-10958 DENA-17-3
## BESC-52   0.000000 0.000000 0.000000 3.741657 0.000000 1.414214  1.414214
## BESC-79   0.000000 0.000000 0.000000 3.741657 0.000000 1.414214  1.414214
## BESC-246  0.000000 0.000000 0.000000 3.741657 0.000000 1.414214  1.414214
## CA-05-06  3.741657 3.741657 3.741657 0.000000 3.741657 3.464102  3.464102
## BESC-460  0.000000 0.000000 0.000000 3.741657 0.000000 1.414214  1.414214
## GW-10958  1.414214 1.414214 1.414214 3.464102 1.414214 0.000000  2.000000
## DENA-17-3 1.414214 1.414214 1.414214 3.464102 1.414214 2.000000  0.000000
```

#### 6.2. Principal Components Analysis

As shown in my publication on Analysis of Molecular Variance (AMOVA) with vegan and BiodiversityR, including a graphical method to identify potential migrants, function **BiodiversityR::caprescale** provides details on different methods of obtaining sums-of-squares from **vegan::capscale** results. With the Euclidean distance, function **vegan::capscale** results in the same configuration as **vegan::rda** that directly calculates a Principal Component Analysis.

```
ind.pca <- caprescale(capscale(Poptri.genind@tab ~ 1,
                              method="euclidean"),
                      verbose=TRUE)

## SSTot obtained from sum of all eigenvalues: 2177.718
## SSTot obtained from sum of all positive eigenvalues: 2177.718
## SSTot reflected by distances among site scores on all axes: 2177.718

ind.pca

## Call: capscale(formula = Poptri.genind@tab ~ 1, method = "euclidean")
##
##                 Inertia Rank
```

```
## Total           2.417
## Unconstrained   2.417    5
## Inertia is mean squared Euclidean distance
## Species scores projected from '@' 'Poptri.genind' 'tab'
##
## Eigenvalues for unconstrained axes:
##   MDS1   MDS2   MDS3   MDS4   MDS5
## 1.0639 0.6884 0.2356 0.2242 0.2049

n.axes <- length(ind.pca$CA$eig)
n.axes

## [1] 5

ind.pca.summary <- summary(ind.pca, scaling=1)

# Relationship between inertia and SSTot
ind.pca$tot.chi * (nrow(Poptri.genind@tab) - 1)

## [1] 2177.718

ind.scores <- summary(ind.pca,
                      axes=n.axes,
                      scaling=1)$sites
```

#### 6.3. Comparison of distance matrices

As expected, the distances among the individuals in PCA / caprescale space correspond to the distances of the Euclidean distance matrix.

```
ind.distB <- as.matrix(vegdist(ind.scores,
                               method="euclidean"))[1:7, 1:7]

ind.distB

##                BESC-52      BESC-79     BESC-246 CA-05-06     BESC-460 GW-10958
## BESC-52   0.000000e+00 1.819315e-13 6.997107e-14 3.741657 1.678682e-14 1.414214
## BESC-79   1.819315e-13 0.000000e+00 1.942472e-13 3.741657 1.783605e-13 1.414214
## BESC-246  6.997107e-14 1.942472e-13 0.000000e+00 3.741657 6.808976e-14 1.414214
## CA-05-06  3.741657e+00 3.741657e+00 3.741657e+00 0.000000 3.741657e+00 3.464102
## BESC-460  1.678682e-14 1.783605e-13 6.808976e-14 3.741657 0.000000e+00 1.414214
## GW-10958  1.414214e+00 1.414214e+00 1.414214e+00 3.464102 1.414214e+00 0.000000
## DENA-17-3 1.414214e+00 1.414214e+00 1.414214e+00 3.464102 1.414214e+00 2.000000
##           DENA-17-3
## BESC-52    1.414214
## BESC-79    1.414214
## BESC-246   1.414214
## CA-05-06   3.464102
## BESC-460   1.414214
## GW-10958   2.000000
## DENA-17-3  0.000000

# Distances calculated earlier in the distance matrix
round(ind.distA,
      digits=6)[1:7, 1:7]

##            BESC-52  BESC-79 BESC-246 CA-05-06 BESC-460 GW-10958 DENA-17-3
## BESC-52   0.000000 0.000000 0.000000 3.741657 0.000000 1.414214  1.414214
## BESC-79   0.000000 0.000000 0.000000 3.741657 0.000000 1.414214  1.414214
## BESC-246  0.000000 0.000000 0.000000 3.741657 0.000000 1.414214  1.414214
```

```
## CA-05-06  3.741657 3.741657 3.741657 0.000000 3.741657 3.464102  3.464102
## BESC-460  0.000000 0.000000 0.000000 3.741657 0.000000 1.414214  1.414214
## GW-10958  1.414214 1.414214 1.414214 3.464102 1.414214 0.000000  2.000000
## DENA-17-3 1.414214 1.414214 1.414214 3.464102 1.414214 2.000000  0.000000

# easier to see the differences among the individuals in ordination space by rounding
round(ind.distB,
      digits=6)[1:7, 1:7]

##            BESC-52  BESC-79 BESC-246 CA-05-06 BESC-460 GW-10958 DENA-17-3
## BESC-52   0.000000 0.000000 0.000000 3.741657 0.000000 1.414214  1.414214
## BESC-79   0.000000 0.000000 0.000000 3.741657 0.000000 1.414214  1.414214
## BESC-246  0.000000 0.000000 0.000000 3.741657 0.000000 1.414214  1.414214
## CA-05-06  3.741657 3.741657 3.741657 0.000000 3.741657 3.464102  3.464102
## BESC-460  0.000000 0.000000 0.000000 3.741657 0.000000 1.414214  1.414214
## GW-10958  1.414214 1.414214 1.414214 3.464102 1.414214 0.000000  2.000000
## DENA-17-3 1.414214 1.414214 1.414214 3.464102 1.414214 2.000000  0.000000

all.equal(target=ind.distA,
          current=ind.distB)

## [1] TRUE
```

#### 6.4. Locations of population centroids

The locations of the centroids are calculate as average of the scores of the individuals on each axis.

```
pop.centroids <- stats::aggregate(ind.scores ~ Poptri.genind@pop, FUN=mean)
pop.centroids <- data.frame(pop.centroids)
rownames(pop.centroids) <- pop.centroids[, 1]
pop.centroids <- pop.centroids[, -1]

pop.distB <- vegdist(pop.centroids, method="euclidean")
as.matrix(pop.distB)[1:7, 1:7]

##            Chilliwack  Columbia      Dean  Harrison  Homathko   Kitimat
## Chilliwack  0.0000000 0.3774089 0.8201005 0.2792185 0.2301599 0.5370614
## Columbia    0.3774089 0.0000000 0.7770705 0.2906766 0.5501403 0.4477654
## Dean        0.8201005 0.7770705 0.0000000 0.9619599 0.9600223 0.7089083
## Harrison    0.2792185 0.2906766 0.9619599 0.0000000 0.4359262 0.5501690
## Homathko    0.2301599 0.5501403 0.9600223 0.4359262 0.0000000 0.7254203
## Kitimat     0.5370614 0.4477654 0.7089083 0.5501690 0.7254203 0.0000000
## Klinaklini  0.7304113 0.9005057 1.4540145 0.6899141 0.6628944 1.0458250
##            Klinaklini
## Chilliwack  0.7304113
## Columbia    0.9005057
## Dean        1.4540145
## Harrison    0.6899141
## Homathko    0.6628944
## Kitimat     1.0458250
## Klinaklini  0.0000000
```

### 7. Comparison between directly calculated distances from the genpop object and the distances among the centroids

It can be seen now that the Euclidean distances among the centroids directly reflect the distances among the populations. The ratio between the two reflects that ratio that was used to standardize the genind@tab slot.

```
pop.distC <- as.matrix(baseline.dist.scaled) / as.matrix(pop.distB)
pop.distC[1:7, 1:7]

##              Chilliwack Columbia Dean Harrison Homathko Kitimat Klinaklini
## Chilliwack          NaN      186  186      186      186     186        186
## Columbia            186      NaN  186      186      186     186        186
## Dean                186      186  NaN      186      186     186        186
## Harrison            186      186  186      NaN      186     186        186
## Homathko            186      186  186      186      NaN     186        186
## Kitimat             186      186  186      186      186     NaN        186
## Klinaklini          186      186  186      186      186     186        NaN

max(rowSums(gen.comm.b))/10

## [1] 186
```

### 8. Graphical comparison

#### 8.1. Ordination of the populations with Principal Components Analysis of the genpop object

```
pca1.balanced <- rda(gen.comm.b ~ 1)
plot1 <- ordiplot(pca1.balanced, display="sites")
text(plot1, what="sites", col="red", cex=0.6)
```

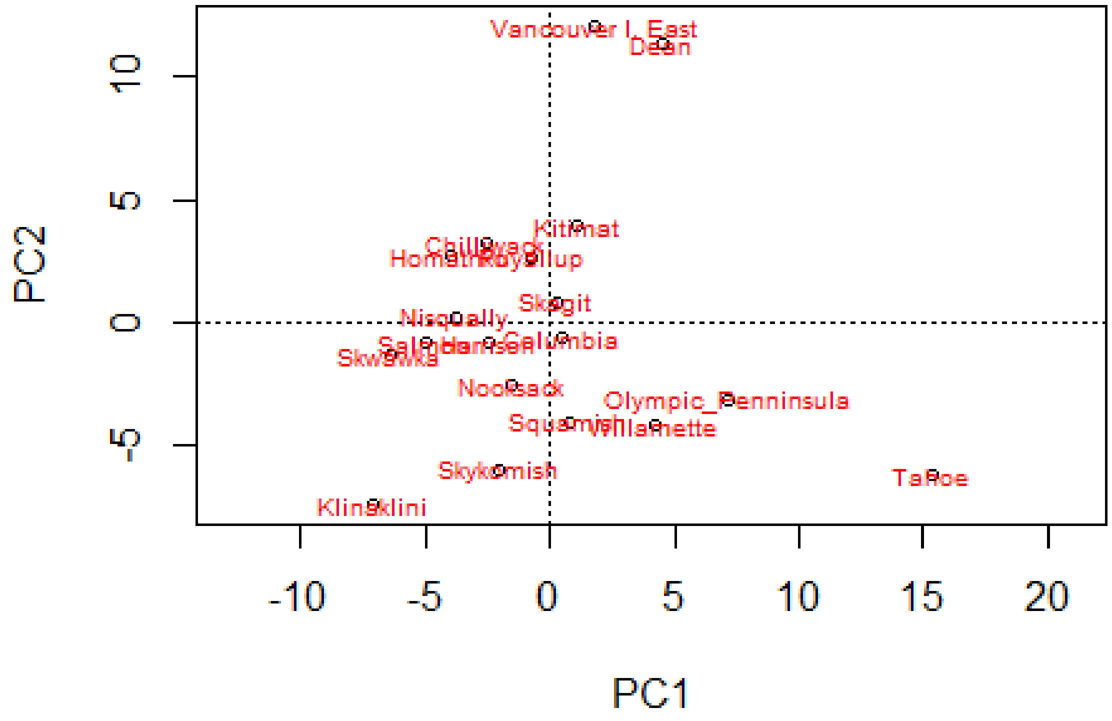

#### 8.2. Ordination of the centroids

```
pca2 <- rda(pop.centroids~1)
plot2 <- ordiplot(pca2, display="sites")
text(plot2, what="sites", col="red", cex=0.6)
```

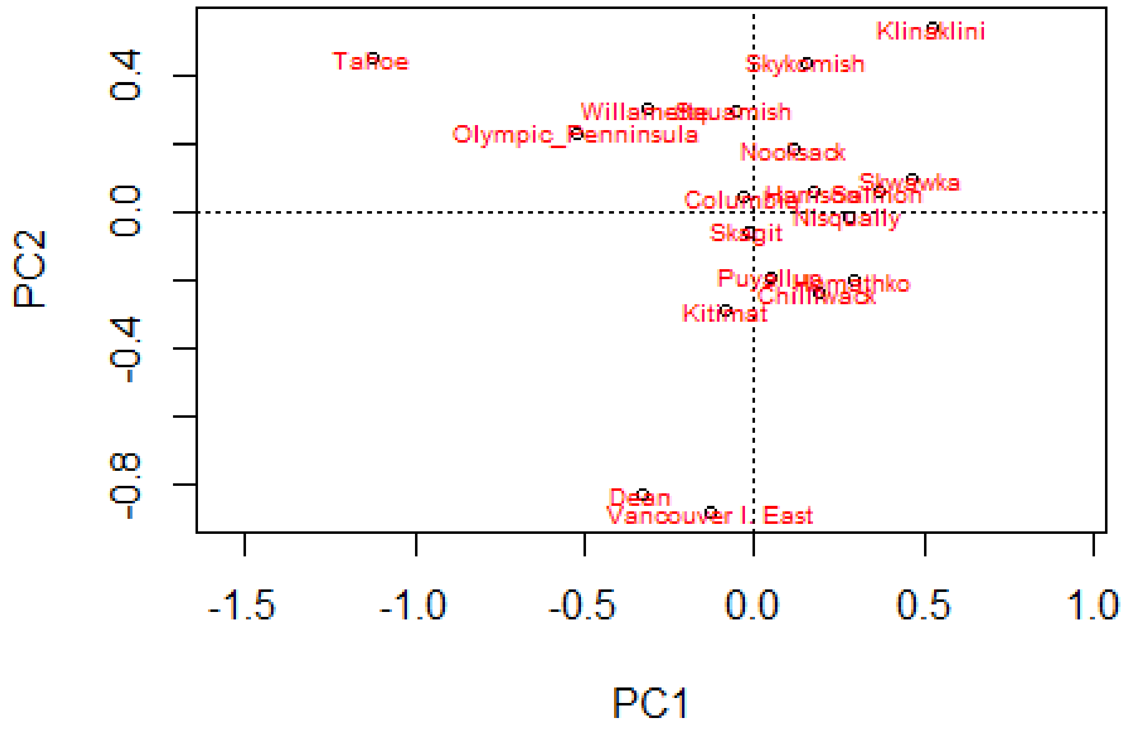

#### 8.3. Flip the scores

As it happens, the locations for the centroids mirror the locations of the first ordination diagram. Obviously, the distances are the same, but it is easier to see that the same configuration is obtained by multiplying positions by -1.

```
pca3 <- pca2
pca3$CA$u[, c(1:2)] <- -1* pca3$CA$u[, c(1,2)]

plot3 <- ordiplot(pca3, display="sites")
text(plot3, what="sites", col="red", cex=0.6)
```

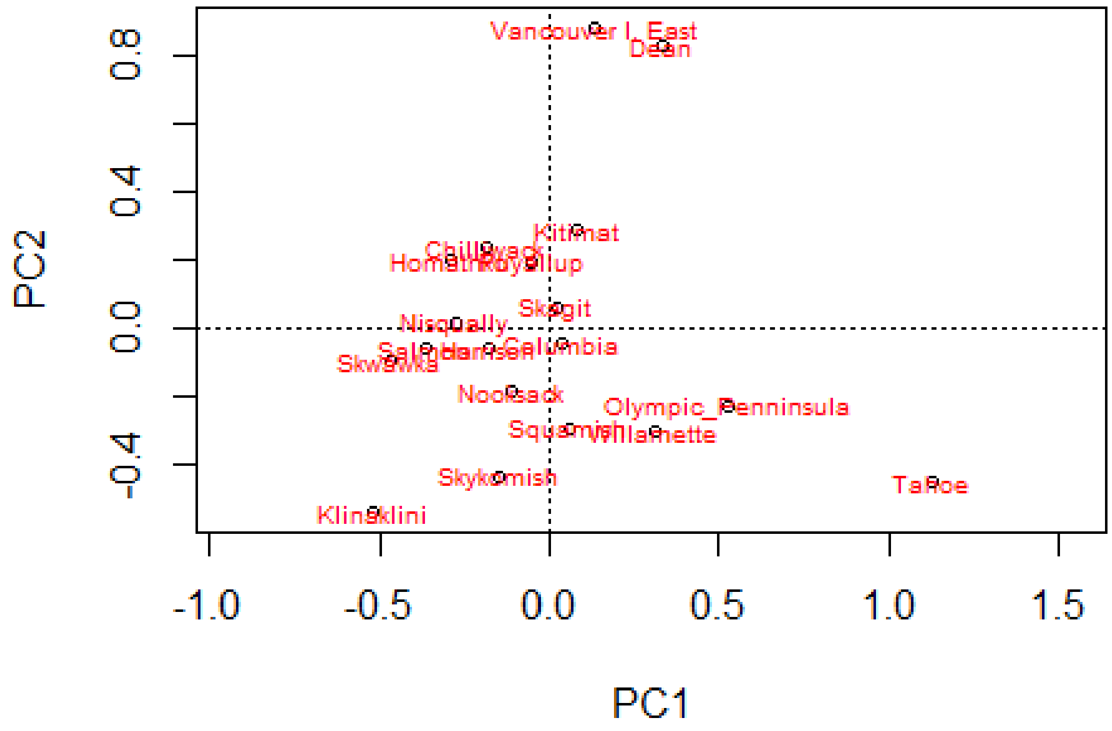

#### 8.4. Comparison between locations

```
scores.plot1 <- data.frame(scores(plot1))
dist.plot1 <- vegdist(rbind(scores.plot1, c(0,0)), method="euc")
arrows.plot1 <- as.matrix(dist.plot1)[, 20]

scores.plot2 <- data.frame(scores(plot2))
dist.plot2 <- vegdist(rbind(scores.plot2, c(0,0)), method="euc")
arrows.plot2 <- as.matrix(dist.plot2)[, 20]

# length of arrow reflect the sums-of-squares
(arrows.plot1/arrows.plot2)^2

##          Chilliwack           Columbia               Dean           Harrison
##                 186                186                186                186
##            Homathko            Kitimat         Klinaklini          Nisqually
##                 186                186                186                186
##            Nooksack Olympic_Penninsula           Puyallup             Salmon
##                 186                186                186                186
##              Skagit            Skwawka          Skykomish           Squamish
##                 186                186                186                186
##               Tahoe  Vancouver I. East         Willamette                 20
##                 186                186                186                NaN
```

### 9. Compare with common genetic distance measures for populations

#### 9.1. Common distances

```
# Euclidean
# method 4: Classical Euclidean distance or Rogers' distance (Euclidean):
Pop.genetic4 <- dist.genpop(Poptri.genpop, method=4)
as.matrix(Pop.genetic4)[1:7, 1:7]

##            Chilliwack   Columbia      Dean   Harrison   Homathko    Kitimat
## Chilliwack 0.00000000 0.03798246 0.1172932 0.04282297 0.02923977 0.07763158
## Columbia   0.03798246 0.00000000 0.1038889 0.04015152 0.06500000 0.06444444
## Dean       0.11729323 0.10388889 0.0000000 0.12662338 0.14126984 0.07500000
## Harrison   0.04282297 0.04015152 0.1266234 0.00000000 0.06212121 0.07500000
## Homathko   0.02923977 0.06500000 0.1412698 0.06212121 0.00000000 0.10277778
## Kitimat    0.07763158 0.06444444 0.0750000 0.07500000 0.10277778 0.00000000
## Klinaklini 0.08223684 0.11916667 0.1732143 0.09204545 0.07638889 0.11250000
##            Klinaklini
## Chilliwack 0.08223684
## Columbia   0.11916667
## Dean       0.17321429
## Harrison   0.09204545
## Homathko   0.07638889
## Kitimat    0.11250000
## Klinaklini 0.00000000

# Nei
Pop.genetic1 <- dist.genpop(Poptri.genpop, method=1)
as.matrix(Pop.genetic1)[1:7, 1:7]

##             Chilliwack    Columbia       Dean    Harrison    Homathko
## Chilliwack 0.000000000 0.003879964 0.01981765 0.002286651 0.001587381
## Columbia   0.003879964 0.000000000 0.01720830 0.002452464 0.009020503
## Dean       0.019817649 0.017208300 0.00000000 0.027470664 0.027347413
## Harrison   0.002286651 0.002452464 0.02747066 0.000000000 0.005762475
## Homathko   0.001587381 0.009020503 0.02734741 0.005762475 0.000000000
## Kitimat    0.008069536 0.004246348 0.01444974 0.008169170 0.015031708
## Klinaklini 0.015846263 0.024091968 0.06340895 0.014063820 0.013018216
##                Kitimat Klinaklini
## Chilliwack 0.008069536 0.01584626
## Columbia   0.004246348 0.02409197
## Dean       0.014449739 0.06340895
## Harrison   0.008169170 0.01406382
## Homathko   0.015031708 0.01301822
## Kitimat    0.000000000 0.03189297
## Klinaklini 0.031892967 0.00000000
```

#### 9.2. Distance-based redundancy analysis for the Rogers' distance

```
cap4 <- caprescale(capscale(Pop.genetic4 ~ 1, method = "euclidean"), verbose=TRUE)

## SSTot obtained from sum of all eigenvalues: 0.1173376
## SSTot obtained from sum of all positive eigenvalues: 0.1173376
## SSTot reflected by distances among site scores on all axes: 0.1173376

cap4

## Call: capscale(formula = Pop.genetic4 ~ 1, method = "euclidean")
##
```

```
##                 Inertia Rank
## Total           0.10708
## Unconstrained   0.11734   10
## Imaginary      -0.01026    8
## Inertia is squared Rodgers distance
##
## Eigenvalues for unconstrained axes:
##    MDS1    MDS2    MDS3    MDS4    MDS5    MDS6    MDS7    MDS8    MDS9   MDS10
## 0.07175 0.01718 0.00998 0.00920 0.00400 0.00199 0.00189 0.00085 0.00040 0.00010

plot4 <- ordiplot(cap4, display="sites")
text(plot4, what="sites", col="red", cex=0.6)
```

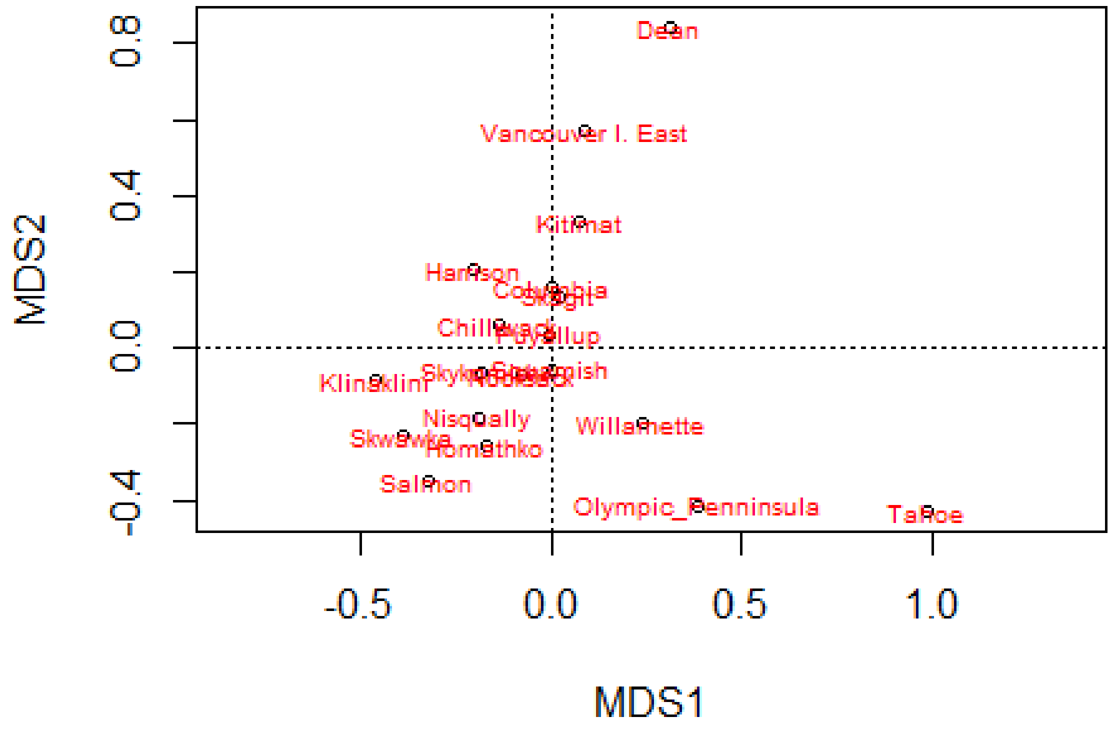

#### 9.3. Procrustes analysis

The procrustes analysis (fun fact: this analysis was named after a figure from Greek mythology that promised guests a perfectly fitting bed for the night, and then stretched the person in case the person was too short and cut of their legs in case they were too tall) shows that most populations occur in the same quadrants.

```
protest(plot4, plot1)

##
## Call:
```

```
## protest(X = plot4, Y = plot1)
##
## Procrustes Sum of Squares (m12 squared):        0.219
## Correlation in a symmetric Procrustes rotation: 0.8837
## Significance:  0.001
##
## Permutation: free
## Number of permutations: 999
```

```
plot5 <- plot(protest(plot2, plot4))
plot5
```

```
## $heads
##                             PC1          PC2
## Chilliwack          0.081622945 -0.103680404
## Columbia           -0.014097079  0.018917125
## Dean               -0.143018530 -0.359889639
## Harrison            0.076087744  0.025578830
## Homathko            0.127200758 -0.085578521
## Kitimat            -0.035348193 -0.124547232
## Klinaklini          0.226089938  0.234665093
## Nisqually           0.120194585 -0.006680199
## Nooksack            0.049745872  0.080578518
## Olympic_Penninsula -0.226502487  0.100197900
## Puyallup            0.021948913 -0.083342967
## Salmon              0.158300579  0.024584125
## Skagit             -0.008402801 -0.025690623
## Skwawka             0.202702758  0.042716854
## Skykomish           0.065911813  0.188586871
## Squamish           -0.024357377  0.128530361
## Tahoe              -0.486879254  0.196409095
## Vancouver I. East  -0.055871554 -0.382534234
## Willamette         -0.135328629  0.131179046
##
## $points
##                             [,1]        [,2]
## Chilliwack          6.062673e-02 -0.02677223
## Columbia           -1.949251e-05 -0.07292452
## Dean               -1.314208e-01 -0.38240859
## Harrison            9.393378e-02 -0.09065576
## Homathko            7.398650e-02  0.11786002
## Kitimat            -3.169061e-02 -0.14920226
## Klinaklini          2.051029e-01  0.04352612
## Nisqually           8.425502e-02  0.08482816
## Nooksack            3.416038e-02  0.03386092
## Olympic_Penninsula -1.785315e-01  0.18209606
## Puyallup            1.030318e-03 -0.01465238
## Salmon              1.413038e-01  0.15944709
## Skagit             -9.093599e-03 -0.06089063
## Skwawka             1.728100e-01  0.10919774
## Skykomish           8.091830e-02  0.02978173
## Squamish           -2.048428e-03  0.02604088
## Tahoe              -4.495500e-01  0.18255896
## Vancouver I. East  -3.513257e-02 -0.25840390
## Willamette         -1.106407e-01  0.08671259
##
## attr(,"class")
## [1] "ordiplot"
```

```
text(plot5, what="points", col="red", cex=0.6)
```

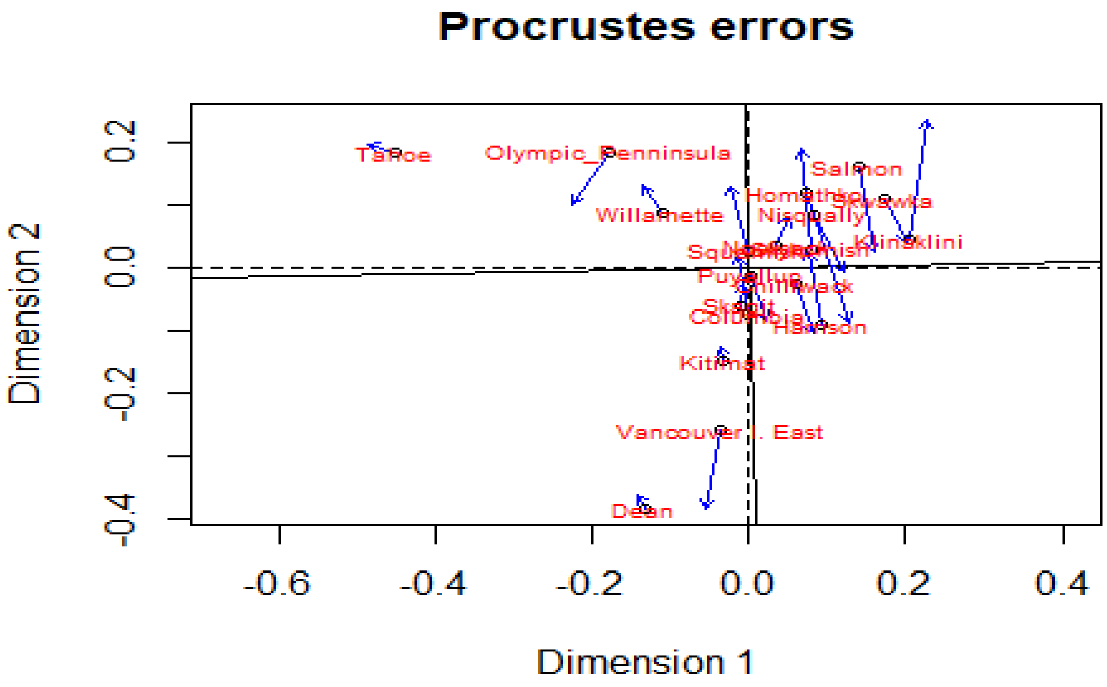

#### 9.4. Distance-based redundancy analysis for the Nei distance

```
cap1 <- caprescale(capscale(Pop.genetic1 ~ 1, method = "euclidean"), verbose=TRUE)

## SSTot obtained from sum of all eigenvalues: 0.02462873
## SSTot obtained from sum of all positive eigenvalues: 0.02462873
## SSTot reflected by distances among site scores on all axes: 0.02462873

cap1

## Call: capscale(formula = Pop.genetic1 ~ 1, method = "euclidean")
##
##                Inertia Rank
## Total           0.01426
## Unconstrained   0.02463    5
## Imaginary      -0.01037   13
## Inertia is squared Nei distance
##
## Eigenvalues for unconstrained axes:
##     MDS1     MDS2     MDS3     MDS4     MDS5
## 0.022773 0.001215 0.000431 0.000179 0.000031

plot5 <- ordiplot(cap1, display="sites")
text(plot5, what="sites", col="red", cex=0.6)
```

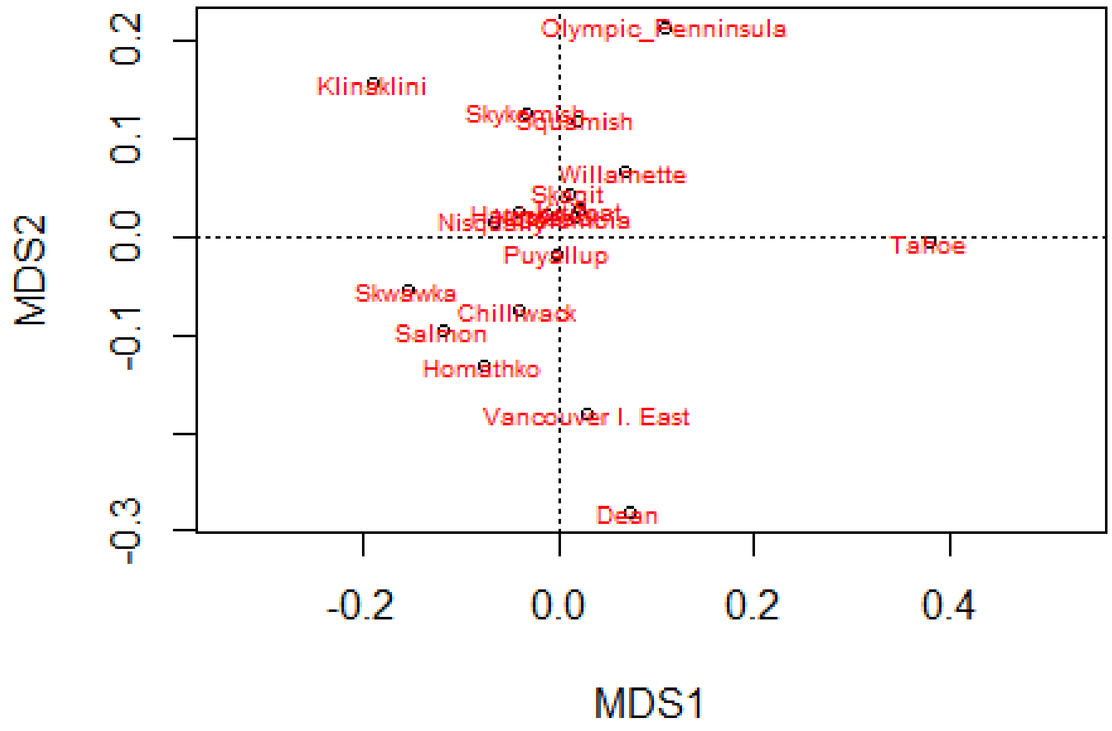

Procrustes analysis

```
protest(plot5, plot1)

## 
## Call:
## protest(X = plot5, Y = plot1) 
## 
## Procrustes Sum of Squares (m12 squared):        0.1985 
## Correlation in a symmetric Procrustes rotation: 0.8953 
## Significance:  0.001 
## 
## Permutation: free
## Number of permutations: 999

plot6 <- plot(protest(plot2, plot5))
plot6

## $heads
##                               PC1          PC2
## Chilliwack           0.081622945 -0.103680404
## Columbia            -0.014097079  0.018917125
## Dean                -0.143018530 -0.359889639
## Harrison             0.076087744  0.025578830
## Homathko             0.127200758 -0.085578521
## Kitimat             -0.035348193 -0.124547232
```

```
## Klinaklini          0.226089938  0.234665093
## Nisqually           0.120194585 -0.006680199
## Nooksack            0.049745872  0.080578518
## Olympic_Penninsula -0.226502487  0.100197900
## Puyallup            0.021948913 -0.083342967
## Salmon              0.158300579  0.024584125
## Skagit             -0.008402801 -0.025690623
## Skwawka             0.202702758  0.042716854
## Skykomish           0.065911813  0.188586871
## Squamish           -0.024357377  0.128530361
## Tahoe              -0.486879254  0.196409095
## Vancouver I. East  -0.055871554 -0.382534234
## Willamette         -0.135328629  0.131179046
##
## $points
##                           [,1]         [,2]
## Chilliwack          0.049295097 -0.093152172
## Columbia           -0.019881824  0.025923304
## Dean               -0.088925258 -0.351405852
## Harrison            0.049971696  0.031500726
## Homathko            0.096105641 -0.164738675
## Kitimat            -0.027412282  0.034848934
## Klinaklini          0.235166719  0.199370734
## Nisqually           0.082664248  0.019075261
## Nooksack            0.015594768  0.027886498
## Olympic_Penninsula -0.137557025  0.266273722
## Puyallup            0.001703421 -0.021547641
## Salmon              0.147974559 -0.118956841
## Skagit             -0.015434898  0.055339193
## Skwawka             0.192461714 -0.065737715
## Skykomish           0.040819584  0.157580043
## Squamish           -0.026451336  0.149137275
## Tahoe              -0.475620376 -0.009516745
## Vancouver I. East  -0.035554222 -0.225138541
## Willamette         -0.084920226  0.083258493
##
## attr(,"class")
## [1] "ordiplot"

text(plot6, what="points", col="red", cex=0.6)
```

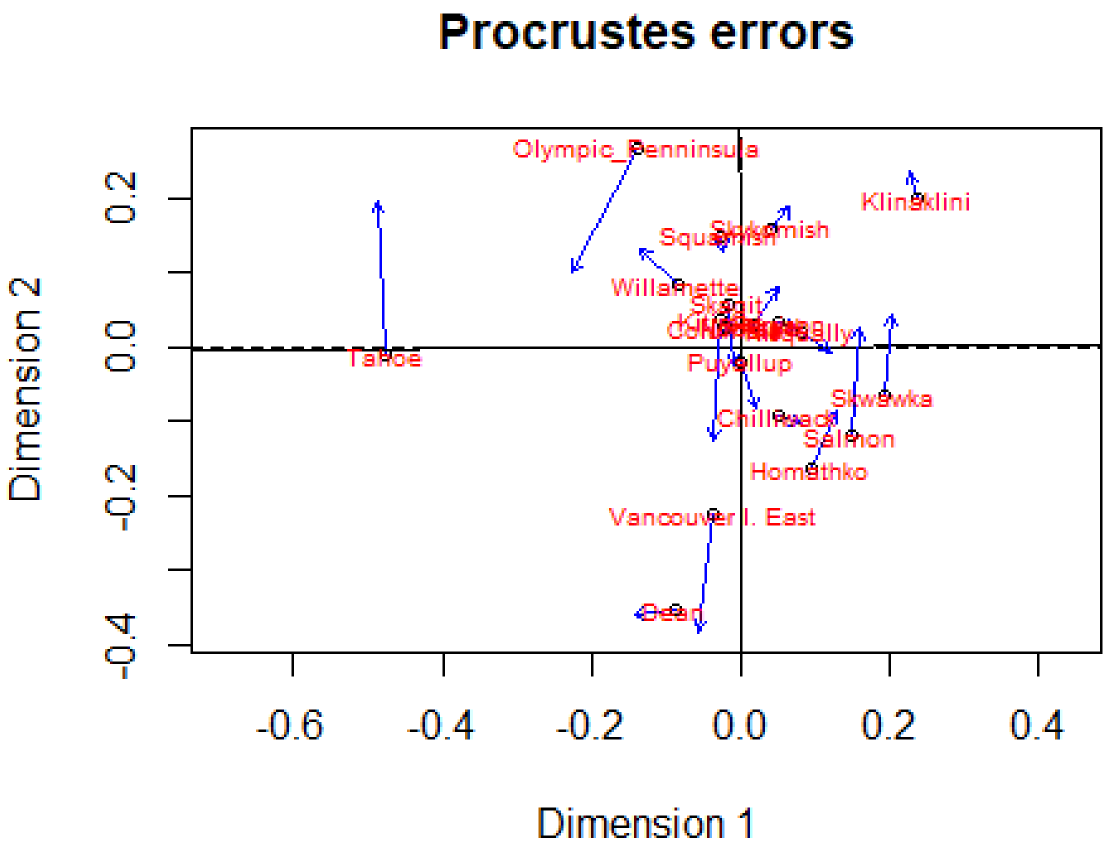
Procrustes errors
Dimension 2
Dimension 1
0.2
0.0
-0.2
-0.4
-0.6
0.4
Olympic_Penninsula
Klinaklini
Squamish
Willamette
Skagit
Puyallup
Tahoe
Chilliwack
Skwawka
Salmon
Homathko
Vancouver I. East
